# Neuronal synchrony and the relation between the BOLD response and the local field potential

**DOI:** 10.1101/083840

**Authors:** Dora Hermes, Mai Nguyen, Jonathan Winawer

## Abstract

The most widespread measures of human brain activity are the blood oxygen level dependent (BOLD) signal and surface field potential. Prior studies report a variety of relationships between these signals. To develop an understanding of how to interpret these signals and the relationship between them, we developed a model of (a) neuronal population responses, and (b) transformations from neuronal responses into the fMRI BOLD signal and electrocorticographic (ECoG) field potential. Rather than seeking a transformation between the two measures directly, this approach interprets each measure with respect to the underlying neuronal population responses. This model accounts for the relationship between BOLD and ECoG data from human visual cortex in V1-V3, with the model predictions and data matching in three ways: Across stimuli, the BOLD amplitude and ECoG broadband power were positively correlated, the BOLD amplitude and alpha power (8-13 Hz) were negatively correlated, and the BOLD amplitude and narrowband gamma power (30-80 Hz) were uncorrelated. The two measures provide complementary information about human brain activity and we infer that features of the field potential that are uncorrelated with BOLD arise largely from changes in synchrony, rather than level, of neuronal activity.

## 1. Introduction

Most measurements of activity in the living human brain arise from the responses of large populations of neurons, spanning the millimeter scale of functional magnetic resonance imaging (fMRI) and electrocorticography (ECoG) to the centimeter scale of electro- and magneto-encephalography (EEG and MEG). Integrating results across methods is challenging because the signals measured by these instruments differ in spatial and temporal sensitivity, and in the manner by which they combine the underlying neuronal population activity [1-3]. Differences in scale can be partially bridged by bringing the measurements into register. For example, EEG and MEG sensor data can be projected to cortical sources subject to constraints from simultaneously recorded fMRI data [4] or from independent fMRI localizers [5], and ECoG electrodes can be aligned to a high resolution anatomical MRI image [6] and compared to the local fMRI signal.

Yet even when electrophysiological and fMRI data are spatially registered, striking differences in the sensitivity to stimulus and task are often observed, indicating differences in how neuronal responses contribute to the measured physiological signals. For example, the fMRI BOLD signal and EEG evoked potentials differ in which brain areas are most sensitive to visual motion (area MT+ with fMRI [7] versus V1 and V3A with EEG [8]). Within the same visual area, fMRI and source-localized EEG evoked potentials can show different effects of task in similar experimental paradigms, such as the effect of spatial attention on the contrast response function (additive in fMRI [9], multiplicative in EEG [10]). Even when the spatial scale of the two signals is approximately matched at acquisition, such as ECoG electrodes and fMRI voxels (both at ~2 mm), systematically different patterns of responses can be obtained, such as compressive spatial summation in fMRI versus nearly linear summation in ECoG steady state potentials (but not ECoG broadband signals) [11]. Such fundamental functional differences cannot be explained by numerical measurement-to-measurement transformations. Rather, these differences must reflect the fact that the measurements are based on different aspects of the neural population response. To explain the differences in measurement modalities requires a computational framework that derives each of these signals from the neuronal responses.

One approach toward developing such a framework has been to measure the BOLD signal and electrophysiological signals simultaneously, or separately but using the same stimulus and task conditions, and to ask how features of the electrophysiological response compare to the BOLD signal. This approach has revealed important patterns, yet after several decades of careful study, some apparent discrepancies remain. A number of studies comparing band-limited power in field potential recordings to the BOLD signal have shown that increases in power between 30 and 100 Hz (gamma band) are more highly correlated with BOLD amplitude than power changes in other bands [12-17]. Yet power changes in this band do not fully account for the BOLD signal: very large power changes can occur in the gamma band without a measurable BOLD signal change [18,19], and power changes in lower frequency bands can be correlated with the BOLD signal independently of power changes in the gamma band [20-23]. It therefore cannot be the case that field potential power in the gamma band is a general predictor of BOLD, even if the two measures are often correlated. Another source of disagreement is that within the gamma band, some reports claim that BOLD is best predicted by synchronous (narrowband) signals [13], and others claim that BOLD is best predicted by asynchronous (broadband) neural signals [11]. Moreover, in some cases, it has been reported that no feature of the local field potential predicts the intrinsic optical imaging signal (closely related to BOLD) as accurately as multiunit spiking activity [24]. Consistent with this claim, a comparison of both motion and contrast response functions measured with single units and with BOLD suggested a tight coupling between BOLD and single unit responses [25-27]. To our knowledge, there is currently no single model linking the electrophysiological and BOLD signals that accounts for the wide range of empirical results.

The numerous studies correlating features of electrophysiological signals with BOLD provide constraints in interpreting the relationship between the two types of signals, yet the approach has not led to a general, computational solution. We argue that one reason that correlation studies have not led to computational solutions is that any particular feature of the field potential could be caused by many possible neuronal population responses. For example, a flat field potential (minimal signal) could arise because there is little activity in the local neuronal population, or it could arise from a pair of neuronal sub-populations responding vigorously but in counterphase, resulting in cancellation in the field potential. The same field potential in the two situations would be accompanied by different levels of metabolic demand and presumably different levels of BOLD signal. Similarly, any particular BOLD measurement could be due to many different patterns of neural activity. For example, stimulation of a neuronal population that inhibits local spiking can cause an elevation in the BOLD signal [28], as can stimulation of an excitatory population that increases the local spike rate [29]. In short, there can be no single transfer function that predicts the BOLD signal from the field potential, because the field potential does not *cause* the BOLD signal; rather, the neuronal activity gives rise to both the field potential and the BOLD signal.

We propose that many of the different claims pertaining to the relationship between BOLD amplitude and features of the field potential can be accounted for by a modeling framework in which BOLD and field potential measurements are predicted from simulated neuronal population activity, rather than by predicting the BOLD signal directly from the field potential. In this paper, we model fMRI and ECoG responses in two stages, one stage in which we simulate activity in a population of neurons, and a second stage in which we model the transformation from the population activity to the instrument measures. By design, the model employs a minimal set of principles governing how the instruments pool neuronal activity, rather than a biophysically detailed description of neuronal and hemodynamic events. This approach enables us to ask whether this minimal set of principles is sufficient to guide simulations of neuronal population activity, such that the parameters of the simulations are fit to ECoG measurements from human visual cortex, and the output of the simulations predicts fMRI BOLD responses in the same regions for the same stimuli.

## 2. Results

*Summary.* We first present an analytic framework to capture basic principles of how the BOLD signal and the field potential pool neuronal signals across a population (2.1). Using this framework, we derive equations for the relationship between each instrument measure (BOLD and LFP) and the underlying neuronal activity, as well as the relationship between the instrument measures. This section shows that synchrony is expected to have a large effect on the LFP signal but not on the BOLD signal. The analytic framework provides a way to derive the instrument measures from neuronal population activity, but it does not specify the neuronal population activity itself. In the next section (2.2), we develop a method for simulating neuronal population time series from a small number of parameterized inputs, and we show how the simulated neuronal activity can be converted to (simulated) LFP and BOLD by applying the equations derived in 2.1. Next, we fit parameters for simulating population neuronal activity using ECoG data from human V1, V2 and V3, and compare the BOLD responses derived from these simulations to measured BOLD responses from V1-V3 (2.3). Finally, we quantify the relationship between simulated BOLD and LFP, and between measured BOLD and ECoG, and show that the same patterns hold for simulation and data (2.4).

### 2.1 Relationship between LFP and BOLD: analytic framework

The fMRI BOLD signal and the local field potential (LFP) measure neuronal population activity in a fundamentally different manner. The goal of this analytic framework is to capture these differences in simple mathematical expressions, and from these expressions derive the relationship between the two instrument measurements. We purposely omit a large number of biophysical details such as cell types, neuronal compartments, the dynamics of blood flow, and so forth, both for tractability and in order to emphasize the basic principles of how different measures integrate neuronal activity. In the sections that follow, we then show that, when coupled to simulated neural responses, the model can account for many important patterns observed in fMRI and ECoG data from human visual cortex.

For this analytic framework, we consider how a population of *n* neurons responds to a stimulus or task during a brief epoch (time 0 to *T*), assumed to be on the order of a second. Each neuron will produce a time varying dendritic current, denoted as *I*_*i*_*(t)* for the *i*th neuron, resulting from the trans-membrane potential. We would like to know how these currents, *I(t)*, relate to the fMRI BOLD signal and to the LFP signal measured by an ECoG electrode.

We assume that the LFP arises primarily from dendritic membrane currents [2]. We ignore output spikes. (Although spikes can influence the LFP [30], it is generally thought that the influence is smaller than synaptic and dendritic currents [2], and including spikes would not change the logic of our arguments.) For the *i*th neuron, the contribution to the LFP is then *α*_*i*_ *× I*_*i*_(*t*). The constant *α*_*i*_ depends on the distance and orientation of the neuron with respect to the electrode, as well as the electrode’s impedance. For simplicity, we assume that each neuron is equidistant from the electrode and has the same orientation, like pyramidal neurons perpendicular to the cortical surface, and therefore its contribution to the electrode measurement is scaled by the same constant,. Because currents add, the LFP time series will sum the contribution from each neuron,

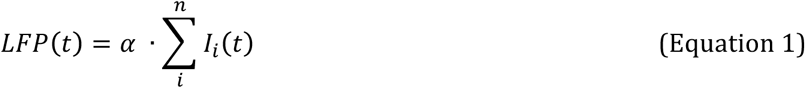

Field potential recordings are usefully summarized as the power (or band-limited power) in the time series [31]. Here we summarize the LFP response within a short time window as the power in the signal summed over the time window T:

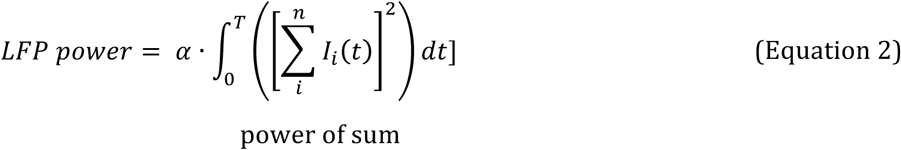

Importantly, Equation 2 is a linear/ nonlinear (L/N) computation, since the LFP power is computed by first summing the signals (L), and then computing the power (N).

The BOLD signal pools neural activity in a fundamentally different manner because it depends on metabolic demand [e.g., for reviews, see 1,32]. The metabolic demand of each neuron will increase if the cell depolarizes (excitation) or hyperpolarizes (inhibition) [28]. Hence the metabolic demand of a neuron is a *nonlinear* function of its membrane potential: either a positive or negative change in voltage relative to resting potential causes a current, thereby resulting in a positive metabolic demand. There are many possible nonlinear functions one could assume to summarize the metabolic demand from the dendritic time series, such as the rectified signal (absolute value) or the power (squared signal). For tractability, we assume the metabolic demand of the *i*th neuron is proportional to the power in the time varying trans-membrane current, integrated over time: *β*_*i*_ *×*(*POWER*(*I*_*i*_(*t*))), or 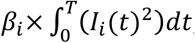, with *β*_*I*_ a scaling constant for the *i*th neuron. (Similar results were obtained if we used the absolute value rather than the power). For the entire population of *n* neurons, we then assume the BOLD signal will sum the metabolic demand of each neuron. For simplicity we use the same scaling constant for each neuron:

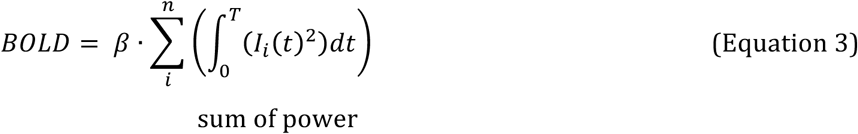

Importantly, Equation 3 is a nonlinear / linear (N/L) computation, since the power is computed first (N) and then the signals are summed (L), opposite to the order of operations for the LFP in Equation 3 (Fig 1) (Personal communication from David J Heeger). In other words, we approximate the BOLD signal as the sum of the power, and LFP as the power of the sum, of the separate neuronal time series. The difference in the order of operations can have a profound effect on the predicted signals, as in the simple example with 2 neurons depicted in Fig 1C **and** 1D. The BOLD signal pooled over the two neurons is the same whether the time series from the two neurons are in phase or out of phase, whereas the LFP power is large when the time series are in phase and small when they are out of phase.

**Fig 1.**
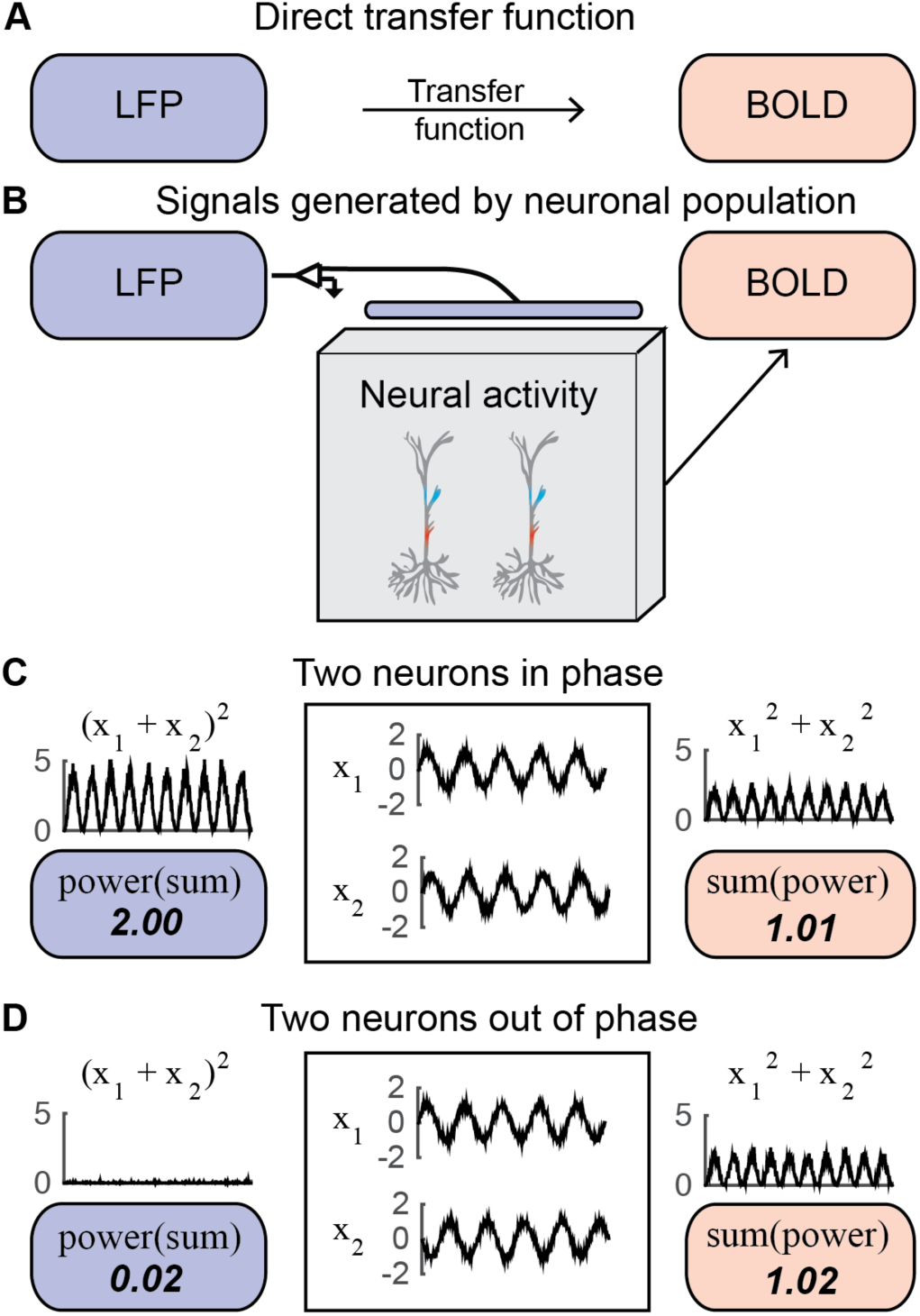
Pooling with different orders of operations can have a large effect on measured brain signals. (A) The approach to directly correlate LFP and BOLD data. (B) Current approach to relate the LFP and BOLD from the same neuronal population activity. (C) In this illustration the membrane potential of two neurons (x_1_ and x_2_) has the shape of a sinusoid with noise, and the sinusoid is in phase between the two neurons. In the simulated electrode measurement, the signals are summed and the power is calculated (POWER(SUM) = 2.00). In the simulated measurement of metabolic demand, the power of each of these neurons is first calculated, and then summed across the neurons (SUM(POWER) = 1.01). Here, the LFP and BOLD are both large. (D) In this illustration the membrane potential of two neurons (x_1_ and x_2_) is the same as in panel (C) except that the two time series are in counterphase. Here, unlike (C), the LFP is nearly 0 and the BOLD signal is large.

These approximations allow us to make predictions about the relation between LFP and BOLD. By theorem, we know that the power of the sum of several time series is exactly equal to the sum of the power of each time series plus the sum of the cross-power between the different time series (Equation 4):

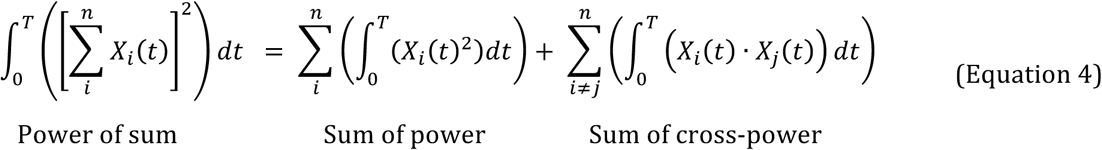

Applying this theorem to Equations 2 and 3 shows the relationship between our models of BOLD and LFP power:

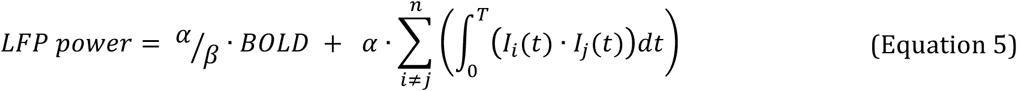

We can now see that the LFP power depends on two quantities, one of which is related to the BOLD signal, and one of which is unrelated to the BOLD signal (Equation 5). The first quantity summarizes the total level of neural activity (summed across neurons), and the second quantity summarizes the relationship between neural time series (the cross-power, similar to covariance). If and when the second term tends to be large compared to the first, then the LFP power will not be closely related to the BOLD signal.

One cannot deduce from first principles whether the first term in Equation 4 (summed power) or the second term (summed cross-power) will dominate. However, the number of elements contributing to the two terms is quite different: For *n* neurons, the first term has *n* numbers (the power in each neuron’s time series), whereas the second term has *n*^*2*^ numbers (all the pairwise cross-powers). Hence if there is any appreciable covariance, then the LFP power will be dominated by the second term, and the correlation with BOLD will be weak (except in cases where the cross-power and power are highly correlated).

To see how these equations translate to quantitative measures of BOLD and LFP, we consider a small neuronal population whose time series conform to a multivariate Gaussian distribution. We assume that each neuron’s time series has the same mean, *m;* the same variance, *σ*^*2*^; and all of the pairwise correlations have the same value, *ρ*:

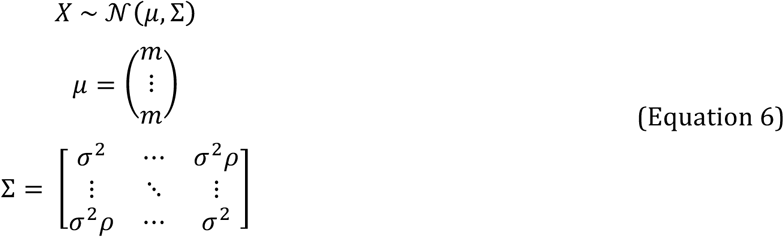

*X* is the population time series, *μ* is the mean of each time series, and ∑ is the covariance matrix. We can now re-write the simulated BOLD signal (the sum of the power) and the LFP (power of the sum) in terms of the parameters of the multivariate Gaussian (and arbitrary scaling factors α, β),

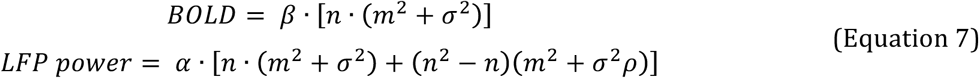

where *n* is the number of neurons. This enables us to visualize how the BOLD signal and the LFP power depend on just 3 values: the variance, correlation, and mean in the neural time series, rather than on all the individual time series (Fig 2). For these neuronal time series, the LFP, modeled as the power of the sum of neuronal time series (panel A), is dominated by the neuronal cross-power (panel C). The BOLD signal, modeled as the sum of the power in the neuronal time series (panel B), makes little contribution to the LFP, except when the correlation between neurons is low (ρ is close to 0); in this case, there is no cross-power, and BOLD and LFP power are correlated.

**Fig 2.**
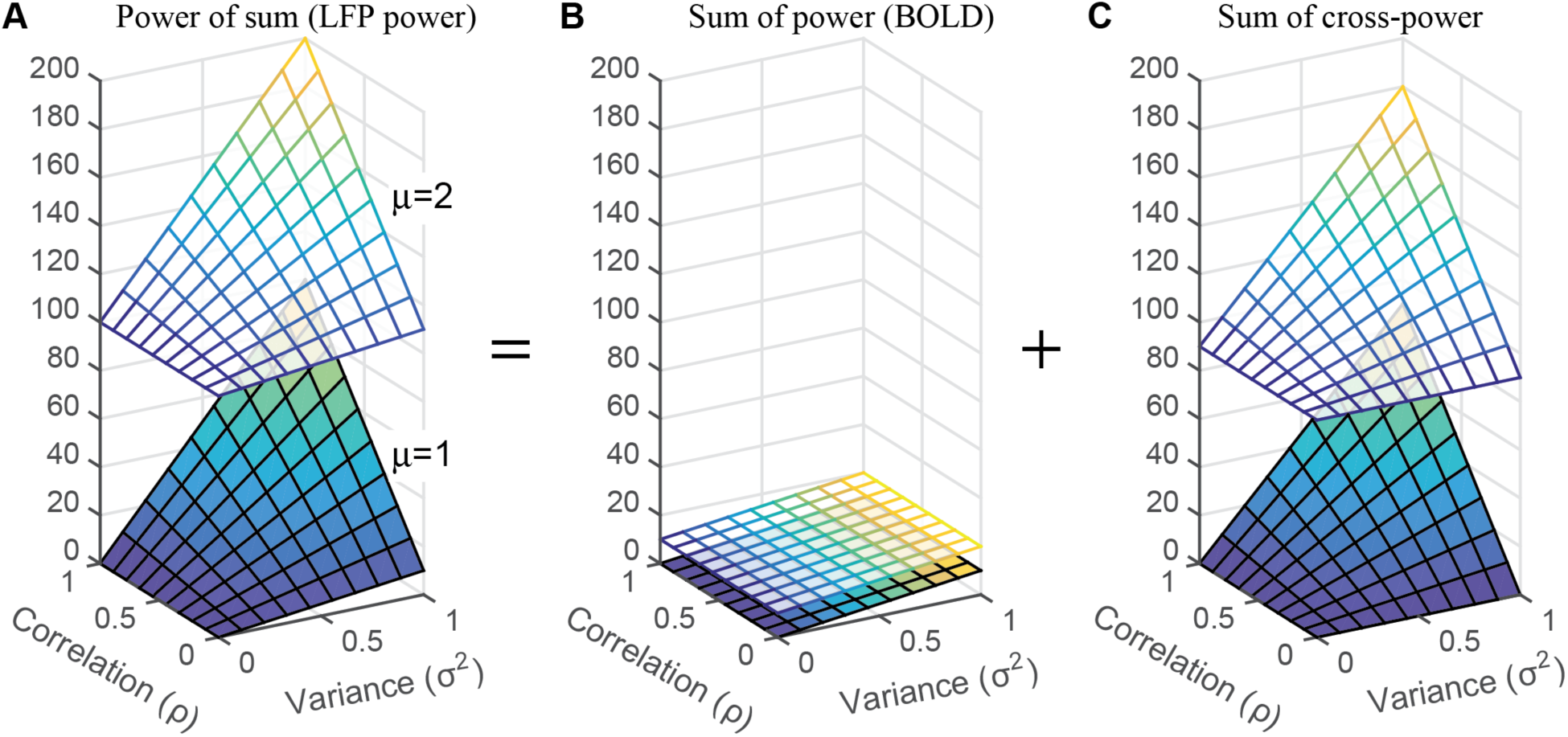
Influence of time series parameters on the power of the sum, the sum of the power and the cross-power. (A) LFP power, computed as the power of the sum of five time series from a multivariate Gaussian distribution (Equation 6). The LFP power is shown as a function of the correlation (ρ), variance (σ), and mean (μ) of the time series (Equation 7). (B) Same as A, except plotting the sum of the power rather than the power of the sum, in order to model the BOLD signal. (C) Same as B but for cross-power. The power of the sum – Panel A – is the sum of the terms in Panels B & C.

### 2.2 Simulating the LFP and BOLD responses

In section 2.1, we proposed formulae to derive instrument measures from neuronal population activity. Here we ask how we might simulate neuronal activity with a small number of parameters. A low dimensional characterization of the population activity is useful since we normally do not have access to the time series of an entire population of neurons. Moreover, a low dimensional representation can lead to better understanding and generalization even when high dimensional data are available [33,34]. After simulating the population activity, we then use the analytic framework from section 2.1 to compute the BOLD and LFP signals. The parameters for the simulations were fit to ECoG recordings from human V1, V2 and V3 [35]. Because there were recordings from multiple electrodes and multiple stimuli, we ran multiple simulations fit to the different ECoG responses. We then used these simulations to predict the BOLD signal and compared these predictions to the measured BOLD signal for the same stimuli and same cortical locations (but in different observers). The steps for simulating the neuronal population data and the derived LFP and BOLD, and for comparing the simulations to empirical data, are summarized in Table 1.

**Table 1.** Summary of analysis steps for simulations and comparison to data.

| Analysis | Figure |
| --- | --- |
| 1. <u>DATA: LFP spectral components.</u> ECoG responses in visual cortex are separated into the sum of 3 spectral components: broadband, narrowband gamma, and narrowband alpha, yielding 3 numbers per electrode per stimulus. | Figure 3 |
| 2. <u>MODEL: Inputs.</u> We propose 3 classes of signals, 'C1', 'C2', and 'C3', as inputs to simulated individual neurons. These are like basis functions, as responses to each stimulus will be modeled with different mixtures of C1, C2 and C3 inputs (Step 5). | Figure 4A |
| 3. <u>MODEL: Neural activity.</u> Neural responses to C1, C2, and C3 inputs are simulated with a population of neurons with leaky integration. | Figure 4B,C |
| 4. <u>MODEL: LFP spectral components.</u> The simulated neuronal activity in the population is converted to simulated LFP. The LFP spectra arising from C1, C2, and C3 inputs approximate the three LFP components observed in ECoG data (step 1 above). | Figure 5 |
| 5. <u>MODEL: Parameters.</u> The model parameters are the levels of C1, C2 & C3 inputs. These are fit to the observed ECoG data for each stimulus and each electrode. This results in 3 numbers per electrode per stimulus. | Figure 6 |
| 6. <u>MODEL: Predictions.</u> These model neural inputs are converted to simulated neuronal activity and then to simulated BOLD responses. | Figure 4D;<br>Figure 7A,B (*x-axis*) |
| 7. <u>MODEL: Accuracy.</u> These simulated BOLD responses are compared with real, measured BOLD. | Figure 7A,B (*y-axis*);<br>Figure 7C |
| 8. <u>MODEL: LFP-BOLD relationship.</u> The simulated BOLD responses are also fit to simulated LFP responses with a regression model. | Figure 8C,D;<br>Figure 9C,D |
| 9. <u>DATA: LFP-BOLD relationship.</u> Real measured BOLD responses are also fit to measured ECoG LFP components using a regression model. | Figure 8A,B;<br>Figure 9A,B |

In the ECoG experiments, there were four grating stimuli of different spatial frequencies, three noise patterns with different power spectra, and one blank stimulus (mean luminance). For each of the 8 stimuli and each of 22 electrodes in V1-V3, we decomposed the measured ECoG responses into three spectral components: broadband, narrowband gamma, and alpha (Fig 3). An important feature of this data set is that the 3 components of the ECoG responses showed different patterns across stimuli [35]: stimuli comprised of noise patterns caused large broadband increases but little to no measureable narrowband gamma response, whereas grating stimuli elicited both broadband increases and narrowband gamma increases. Gratings and noise stimuli both resulted in decreases in alpha power compared to baseline (also see Fig S1). Had all three responses been tightly correlated with each other, it would not be possible to infer how each relates separately to the BOLD signal.

**Fig 3.**
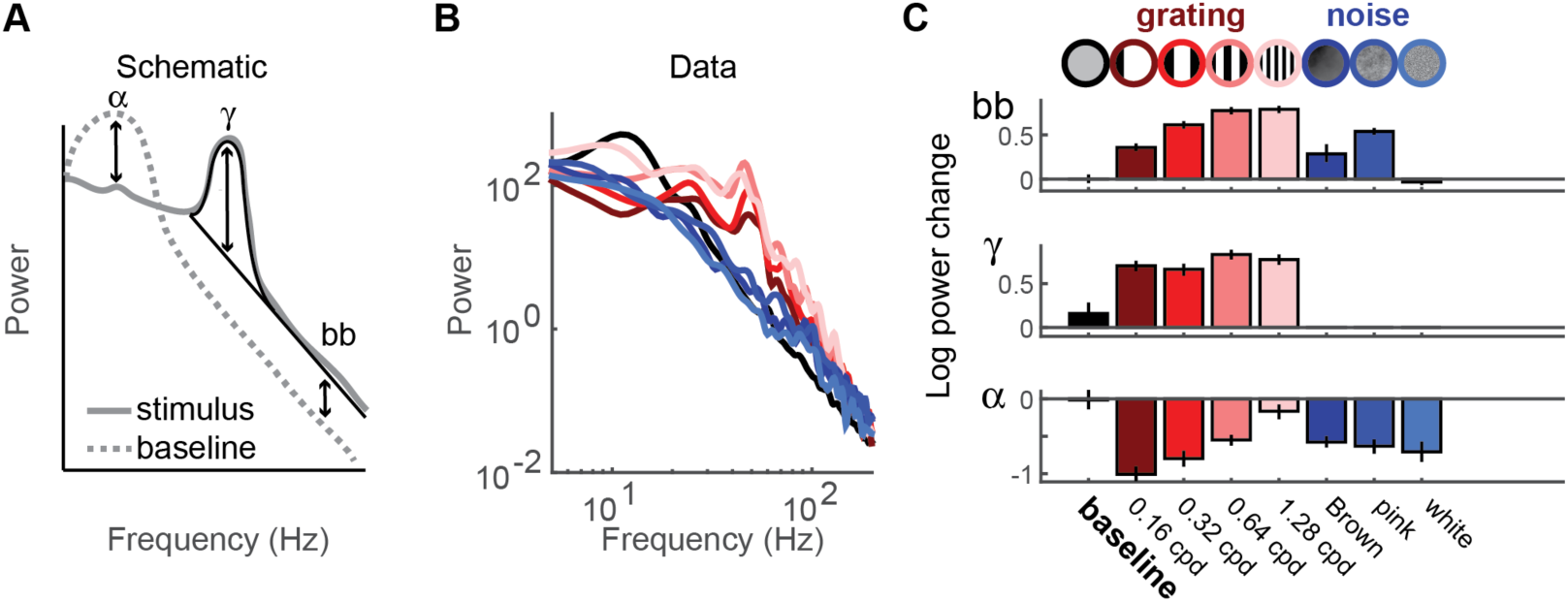
Decomposing ECoG data into 3 summary components. (A) Schematic to show summary metrics derived from ECoG spectra: broadband power elevation (bb), narrowband gamma (γ) and alpha (α). Broadband was calculated by the increase in a 1/f^n^ signal, gamma was calculated by fitting a Gaussian on top of the 1/f^n^ line, and alpha was calculated as the difference from baseline in the alpha-frequency-range. (B) Power spectrum for one example electrode during a blank stimulus (black), gratings (red) and noise patterns (blue). (C) From the power spectrum, changes in broadband, gamma and alpha were calculated. These values were bootstrapped 100 times across trials. Error bars represent 68% confidence intervals.

#### 2.2.1 Simulations of BOLD and LFP responses from neuronal population activity

Cortical neurons receive a large number of inputs from diverse cell types. For our low-dimensional parameterization of the population activity, we assumed that each neuron received a mixture of three types of inputs (Fig 4A). These 3 inputs, following summation and leaky integration, produce the three spectral components observed in the ECoG data. Input 1 approximated Poisson-like spike arrivals (*C*^*1*^, ‘Broadband’), and had a mean above 0 (excitatory). Input 2 was a high frequency oscillation, peaked between 40 and 60 Hz, coordinated between neurons (*C*^*2*^, ‘Gamma’), with a mean of 0. Input 3 was a low frequency signal peaked between 8 and 12 Hz that was inhibitory: i.e. the mean was below 0 (*C*^*3*^, ‘Alpha’).

**Fig 4.**
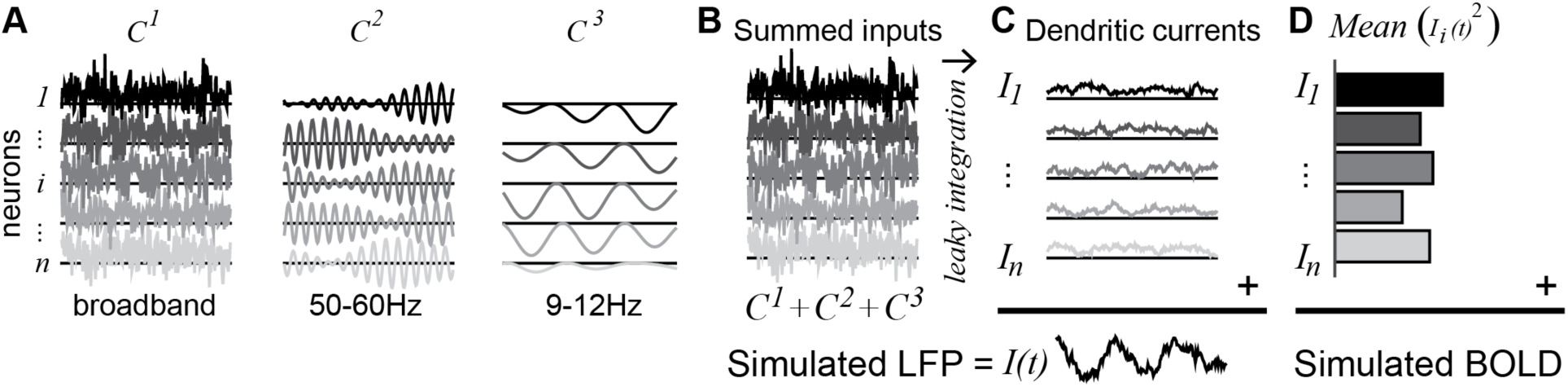
Simulated LFP and BOLD. (A) Three different inputs to each neuron were simulated: a broadband, random input with a small positive offset (*C*^*1*^), an oscillatory input with a time-scale of 40-60Hz (*C*^*2*^), and a negative input with a time-scale of 10 Hz (*C*^*3*^). (B) The three inputs (*C*^*1*^*, C*^*2*^*, C*^*3*^) were summed in each neuron to produce the total input to the neuron. (C) The total input was passed through a leaky integrator to produce the dendritic dipole current (*I*_*i*_). The LFP was simulated by summing the dendritic currents. (D) The BOLD signal was simulated by taking the power of the dendritic current for each neuron and then summing across neurons.

For each simulated neuron *i*, the total input *C* on each trial is the sum of these three signals (Fig 4B):

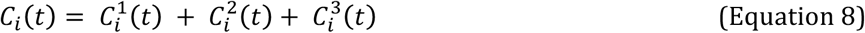

We then passed the summed input in each neuron through a leaky integrator to produce the time-varying dendritic current for that neuron (*I*_*i*_, Fig 4C):

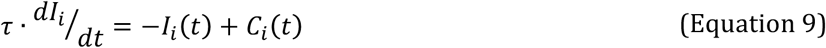

The membrane time constant, τ reflects the time dependence of the trans-membrane current [36,37]. In total, we modeled a population of 200 neurons, each of which produced a one-second time series on each trial. From the neuronal population simulations, we computed the LFP and BOLD signals according to the equations above (section “Relationship between LFP and BOLD: analytic framework”). In brief, the LFP was computed by summing the trans-membrane current across neurons (Equation 1, Fig 4C), and the BOLD signal was computed by summing the power across neurons (Equation 3, Fig 4D). The LFP was used as training data (to fit the parameters of the inputs) and the BOLD was used as test data (to test the accuracy of the model).

#### 2.2.2 Simulation inputs produce three effects in the LFP power spectrum

Below we explain how the time series was generated for each of the three types of inputs and what kind of effect a change in each input has on the power spectrum.

*Broadband input (C*^*1*^). Input *C*^1^ was Gaussian white noise with a small positive bias. The Gaussian white noise approximates Poisson distributed spike arrivals, each of which produces a small positive or negative conductance change, corresponding to excitatory or inhibitory post-synaptic potentials. The small positive bias reflects the assumption of more excitatory than inhibitory synaptic currents, causing a net depolarization. Gaussian white noise was used rather than Poisson distributed synaptic inputs for computational efficiency, but the pattern of results is similar for Poisson or Gaussian distributions. For purposes of simulations, we defined a high value of *C*^1^ as high variance in the Gaussian distribution, and low values of *C*^1^ as low variance. This mimics the effect of high versus low rates of spike arrivals. The time series for the 200 neurons was generated from a distribution with 0 correlation for all pairs of neurons. Because the variance differed across simulations and the correlations were always 0, the possible C^1^ values span a vertical slice of the plots in Fig 2 (*ρ*-axis = 0). This white noise input, after passing through leaky integration, results in an output whose power spectral density declines with temporal frequency. When this input increases (higher variance), the result is a broadband elevation in power [37] (Fig 5A). Such broadband power elevations can be observed in the local field potential [38] as well as intracellular membrane potentials of single neurons in awake macaque [39].

**Fig 5.**
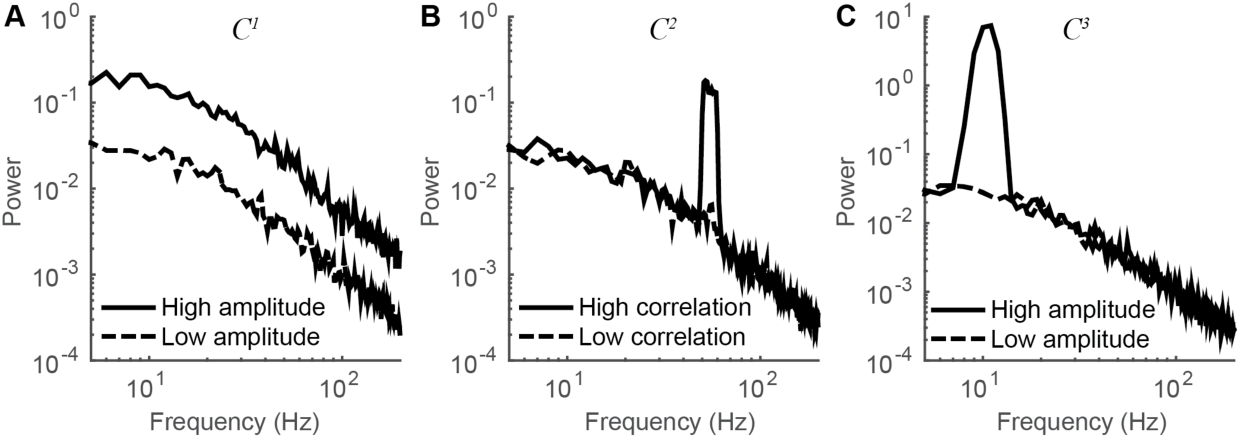
Effect of varying simulated neural inputs on output spectra. The effect of manipulating one of the three neural inputs used in the simulations produced different effects in the spectral power of the LFP of 200 neurons. (A) For *C*^*1*^ (broadband), a high amplitude results in a broadband power elevation, with no narrow peaks in the spectrum. (B) For *C*^*2*^ (gamma), a high correlation results in a narrowband gamma power elevation, with no broadband elevation or change in alpha power. (C) For *C*^*3*^ (alpha), a high amplitude input results in a narrowband power elevation in the alpha band, with no change in broadband power or narrowband gamma power. For each spectrum in each plot, 10 simulated trials were run. The plotted spectra are averaged across the 10 trials, and are computed from *I(t)*, the time series after leaky integration of the inputs.

G*amma input.(C*^*2*^) Input *C*^2^consisted of band pass noise (40-60 Hz), with fixed amplitude on all trials, and with coherence across neurons that varied between trials. This input approximates the signals giving rise to narrowband gamma oscillations. Across different conditions, we varied the correlation between neurons of *C*^2^ rather than the amplitude for individual neurons, which was fixed. This corresponds to a slice in the plots in Fig 2 such that the variance axis is fixed at a non-zero value. The motivation for this comes from empirical observations that large gamma oscillations in the LFP tend to reflect increased coherence between neurons [40,41]. This is opposite to the broadband input (*C*^1^), for which we varied the amplitude (variance) in individual neurons across trials, rather than the synchrony between neurons. Narrowband gamma oscillations with a peak between 30 and 80 Hz can be observed in the local field potential [42,43], as well as in the membrane potential of individual pyramidal neurons [44]. When we increase the correlation of *C*^2^ in our simulations, the result is an increase in the amplitude of the LFP in the gamma band (Fig 5B), much like narrowband gamma signals observed in microelectrode recordings [45] and human ECoG [35].

*Alpha input (C*^3^). The alpha input consisted of inhibitory oscillations at approximately 10 Hz, with fixed correlation between neurons, and varying amplitude across conditions. This corresponds to a slice in the plots in Fig 2 in which the *ρ-*axis is fixed at a non-zero value. The oscillations were inhibitory, i.e. the mean was below 0 (compare *C*^3^ versus *C*^1^ and *C*^2^ in Fig 4). Because *C*^3^ was inhibitory, it resulted in less depolarization (or hyperpolarization in extreme cases), opposite the effect of *C*^1^, which resulted in depolarization. This input approximates the signals giving rise to alpha oscillations (Fig 5C). Pyramidal neurons in visual cortex have been hypothesized to receive periodic inhibition, with pulses arriving at approximately 10 Hz [46,47]. Individual neurons in visual cortex can indeed show subthreshold membrane oscillations at frequencies around 10 Hz [48].

### 2.3 Fitting the simulation parameters to ECoG responses and predicting BOLD data

The simulations were structured to approximate the experimental design and the results of our ECoG experiments. To match the design of our ECoG experiments, a simulated experiment consisted of 240 trials of 1 sec long (30 repeats of 8 conditions). The LFP time series were transformed to power spectra, which were averaged across the 30 repeated trials of the same condition. The simulation parameters – i.e., the level of the three inputs, *C*^*1*^, *C*^*2*^, and *C*^*3*^ – were fit to the measured ECoG summary metrics (broadband, gamma and alpha) for each of the 8 conditions for a particular electrode (Fig 6). To verify the validity of this procedure, we asked whether the simulations using the fitted parameters produce simulated spectra which, when analyzed like the ECoG spectra, reproduce the original values of broadband, gamma, and alpha. In other words, do we close the loop from measured spectral components (broadband, gamma and alpha) to inferred input parameters (*C*^*1*^, *C*^*2*^, *C*^*3*^) to simulated population activity, to simulated spectral components (broadband, gamma and alpha)? The original values are not reproduced exactly because the simulation are stochastic, but overall the original broadband, gamma, and alpha values are recovered with high accuracy (Fig S9).

**Fig 6.**
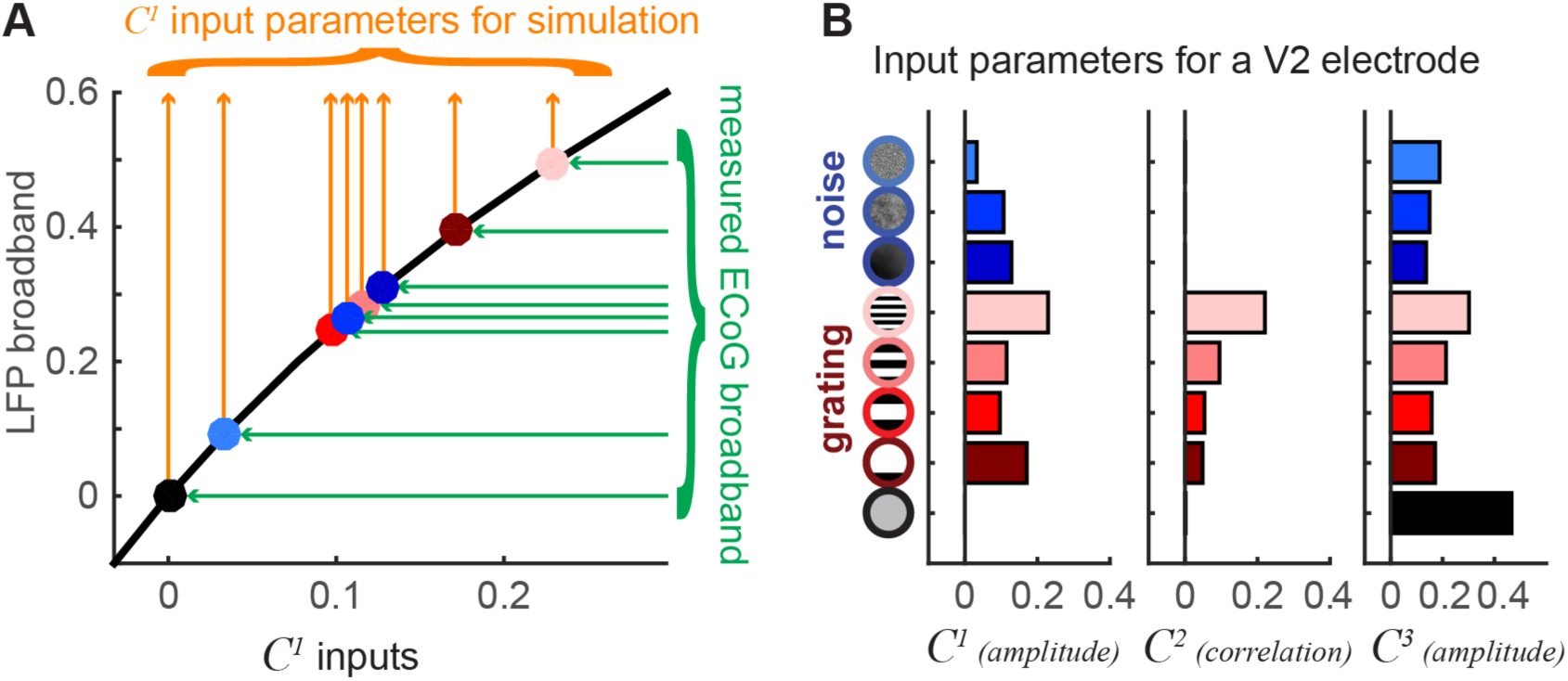
Parameter fits for simulations. (A) A function was fit between the *C*^*1*^ input and the simulated broadband output (black line). Inverting this function allows us to take the measured ECoG broadband values (green arrows to dots colored by stimulus condition), and convert these into estimates of the *C*^*1*^ input levels (orange arrows). Because the simulations contain noise, the predicted broadband need not match the measured broadband exactly; however, the agreement is close, as shown in Fig S9. The parameters for *C*^*2*^ and *C*^*3*^ were fit similarly. (B) The *C*^*1*^, *C*^*2*^ and *C*^*3*^ parameters for simulation of a responses in a V2 electrode for 9 stimuli (8 contrast patterns plus a blank condition). These parameter fits were made using the full ECoG data set (for this electrode), so there are no error bars on the inputs parameters.

As described above, the fitting of the parameters for *C*^*1*^, *C*^2^ and *C*^3^ was constrained by the assumptions that for *C*^1^, the correlation between neurons was zero (and the amplitude was varied for fitting); for *C*^2^ the amplitude was fixed at a non-zero value (and the correlation was varied for fitting); and for *C*^*3*^, the correlation was fixed at a non-zero value (and the amplitude was varied for fitting). Results from alternative models with different constraints show poorer fits, and are described briefly below and more extensively in the Supplement.

Importantly, the parameter fits did not take into account the measured BOLD responses. Hence the simulations provided a test: if the input parameters were chosen to produce outputs that match the measured ECoG responses (training data), does the simulated BOLD signal accurately predict the measured BOLD signal (test data)? We measured BOLD responses in 4 healthy subjects to the same visual stimuli as used in ECoG (subjects are different from the ECoG subjects), and extracted the signal from regions of interest in visual cortex matched to the previously recorded ECoG electrode locations (Supplemental Fig S2 **and** S3). For an example V1 site, the predicted BOLD signal accurately matched the measured BOLD, with 89% of the variance in the measured BOLD explained by the prediction, as quantified by *R*^*2*^, the coefficient of determination (Fig 7A). Across V1 sites, the predicted BOLD signal from the simulations accounted for a median of 80% of the variance in the measured data (Fig 7C). For an example V2 site, the predicted BOLD signal also matched the measured BOLD signal (*R*^*2*^ = 0.74, Fig 7B). Across V2/V3 sites the simulations explained a median of 40% of the variance in the data. The explained variance in V2/V3 is substantial but lower than in V1. One likely reason for the higher variance explained in V1 is that for the particular stimuli used in these experiments (gratings and noise patterns), the BOLD response reliability was higher in V1. For example, the median *R*^*2*^ computed by using half the BOLD data as a predictor for the other half (split half by subjects) was 86% for V1 and 63% for V2/V3. Similarly, the stimulus evoked BOLD responses in V1 were larger than in V2 and V3, with more stimulus-related variance to explain: a mean of 1.84% signal change in V1 versus 1.18% in V2 and 0.81% in V3 (Supplemental Fig S4). It is possible that a stimulus set more tailored to extrastriate areas, such as textures or more naturalistic scenes, would have evoked more reliable responses in extrastriate cortex.

**Fig 7.**
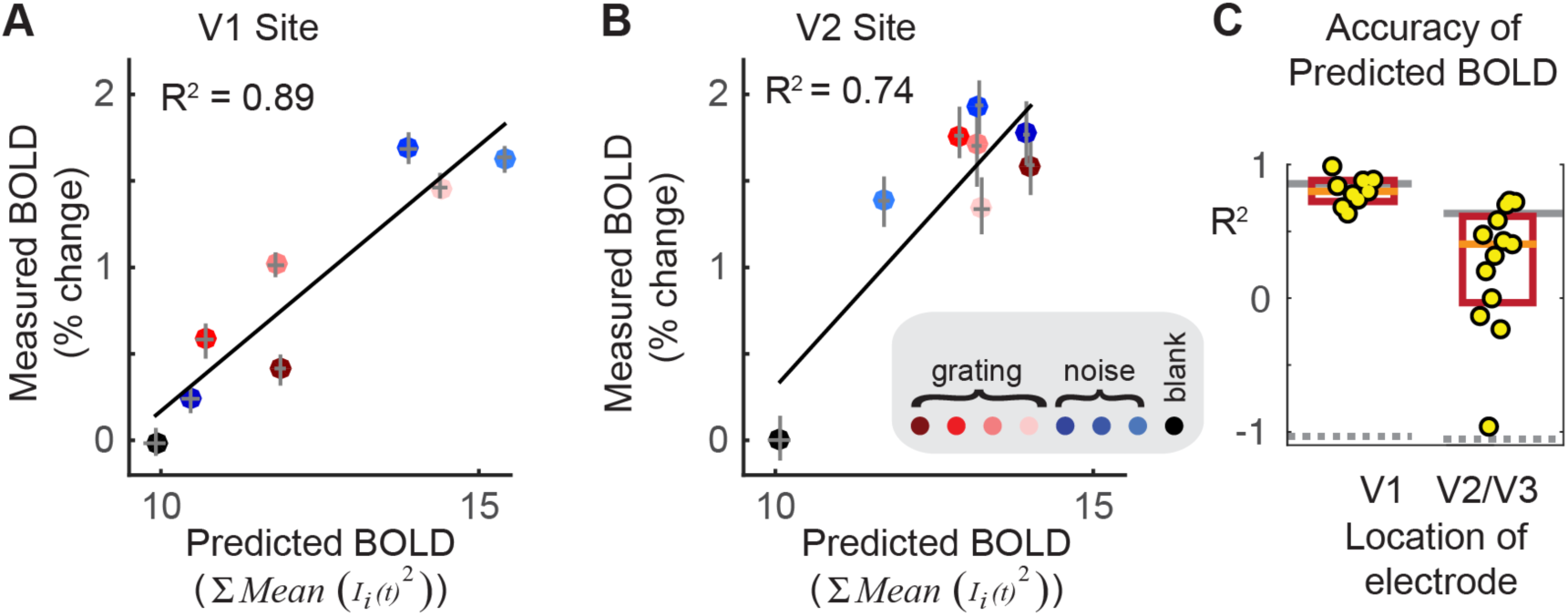
Accuracy of predicted BOLD signals from simulated neuronal activity. (A) Simulated BOLD (x-axis) versus measured BOLD (y-axis) for a V1 site. Each color corresponds to one stimulus condition (red dots, grating patterns; blue dots, noise patterns; black dot, uniform stimulus, or blank). Error bars indicate 68% confidence intervals, bootstrapped 100 times over 30 trials per stimulus for simulation and over repeated scans for BOLD data. (B) Same as A, but for a V2 site. (C) Accuracy of BOLD predictions for all V1 and V2/V3 sites. Each site is indicated by a yellow dot. The orange lines show the medians and the red boxes the 0.25 and 0.75 quantiles. The thin gray solid lines show the BOLD data-to-data reliability, and the gray dashed lines show the accuracy when the BOLD data and trial conditions are shuffled in the training data set. Accuracy is quantified as the coefficient of determination after subtracting the mean from the data and the predictions, and dividing each variable by its vector length. Because the simulations were fit to ECoG data and tested on BOLD data, the predictions are cross-validated, and the coefficient of determination spans (-∞, 1]. A value of -1 is expected when the data and predictions are unrelated and have equal variance, as in the case of the shuffled control analysis.

For each of the 22 simulations, the three input parameters *C*^*1*^, *C*^*2*^, and *C*^*3*^ defining each of the 8 stimulus conditions were fit to produce the LFP data from the corresponding ECoG electrode. By design, the *C*^*1*^ (broadband) and *C*^*3*^ (alpha) inputs were fit to ECoG data by varying the *level* per neuron, whereas *C*^*2*^ was fit to data by varying the *correlation* across neurons. In principle, for any of the three inputs, the ECoG data could have been fit by varying either the level per neuron or correlation across neurons. For completeness, we tested all 8 combinations of models (Supplemental Fig S7). The most accurate model, quantified as the *R*^*2*^ between the measured BOLD and the simulated BOLD (median across the sites in V1 or in V2/V3), was the simulation type used in the main text, in which *C*^*1*^ and *C*^*3*^ varied in the level per neuron and *C*^*2*^ varied in the correlation across neurons. Models in which the broadband correlation rather than level was used to fit the ECoG broadband power were much less accurate. The models in which the gamma LFP power was fit by modulating the level rather than the correlation in the simulated population caused a small drop in *R*^*2*^.

### 2.4 Relation between ECoG and BOLD in simulation and data

The previous analysis showed that when simulations were fit to ECoG data, the simulated BOLD response predicted the measured BOLD response. Here we used regression analysis to assess how the simulated LFP predicted the simulated BOLD, and how the measured LFP predicted the measured BOLD.

#### 2.4.1 Examples of the relation between the BOLD amplitude and LFP features

We first consider an example V1 site (Fig 8A – same site plotted in Fig 7A). The BOLD amplitude for the different stimuli was accurately predicted by the broadband response (*R*^*2*^ = 0.85), but not by the narrowband gamma or alpha power (*R*^*2*^ = -0.06, *R*^*2*^ = -0.07, respectively). Hence, in V1, only broadband power was a good predictor of BOLD amplitude (indicated by the black outlines in Fig. 8). Because there were only 8 data points to fit in each of the 3 correlations, we used cross-validation to avoid overfitting: Linear regression was used to fit the BOLD signal to the ECoG measure in separate halves of the data, and the *R*^*2*^ was computed from the left-out half of each data set.

**Fig 8.**
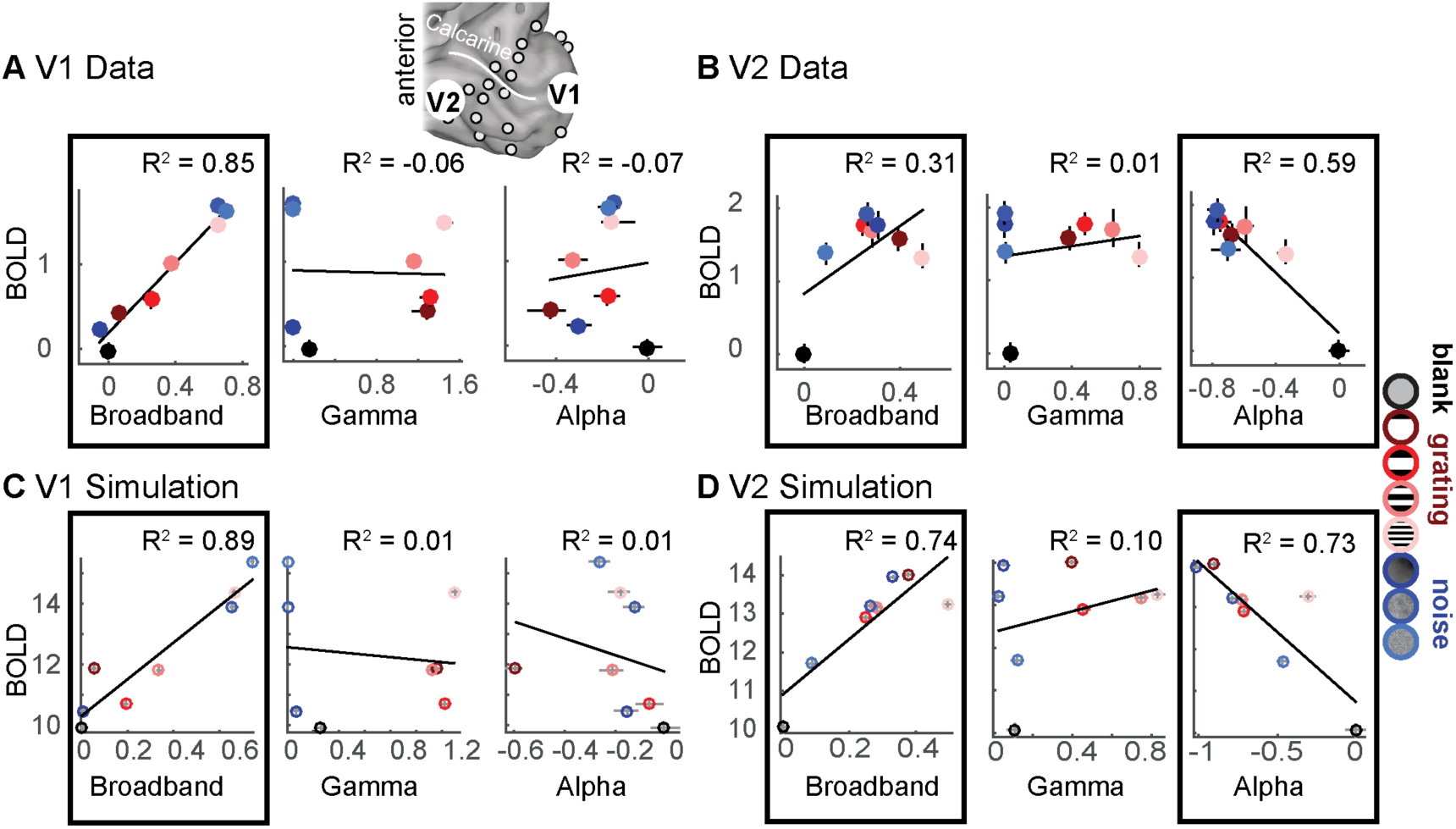
Accuracy of predicted BOLD signals from ECoG components. The correlation between ECoG and BOLD was calculated for a V1 site and a V2 site. The locations of one sample electrode in V1 and one in V2 are indicated by the enlarged white discs on the cortical surface for subject 1. (A) In a foveal V1 site, the broadband ECoG amplitude accurately predicted the BOLD signal (left). Error bars show 68% confidence intervals across bootstraps. Narrowband gamma power (center) and alpha power (right) was uncorrelated with BOLD. (B) In a V2 site, the broadband ECoG was weakly correlated with BOLD (left). Narrowband gamma did not predict BOLD (middle). Alpha was negatively correlated with BOLD (right). Scatter plots for all other V1 and V2/V3 sites are shown in Fig S5. (C-D) Same as A,B, but for simulated neuronal population data from fit to the V1 and V2 ECoG data. For all panels: data points are the bootstrapped median across 30 trials per stimulus (ECoG) and across scans (BOLD). The trend lines are least square fits to the 8 data points plotted. The *R*^*2*^ values are the coefficient of determination computed by cross-validation, with a regression fit to half the data and evaluated on the other half. The black outlines indicate the regressions that show reliable predictors of the BOLD signal – broadband in V1; broadband and alpha in V2/V3.

A different pattern was found in the V2/V3 data. For an example V2 site (Fig 8B – same site plotted in Fig 7B), the broadband power did not predict the BOLD signal as accurately as it did for the V1 site (*R*^*2*^ = 0.31), whereas the alpha power predicted the BOLD signal more accurately than it did for the V1 site (*R*^*2*^ = 0.59). Gamma power did not predict the BOLD signal, similar to the V1 data.

For the simulation fit to the ECoG data from the example V1 electrode, the relation between the BOLD signal and the LFP (Fig 8C) was similar to the measured V1 data: The LFP broadband response predicted the BOLD signal (*R*^*2*^ = 0.89), whereas the power of narrowband gamma oscillations and alpha oscillations did not (*R*^*2*^ = 0.01). In this simulation, similar to the data, the LFP broadband response was the only good predictor of the BOLD signal. Hence the data and the simulation match in that they both show that some features of the LFP are good predictors of BOLD (broadband) and some are poor predictors (alpha and gamma).

For the simulation fit to the ECoG data from the example V2 electrode, the relation between the BOLD signal and the LFP (Fig 8D) was similar to the measured V2 data. First, the broadband LFP was again a good predictor of BOLD, and gamma power was again a poor predictor. Second, the power of alpha oscillations was strongly negatively correlated with BOLD (*R*^*2*^ = 0.73).

In the simulations, the correlation between broadband and BOLD is higher for V1 than for V2, and the correlation between alpha and BOLD is higher for V2 than for V1. These differences were not due to a difference in the types of inputs (C^1^, C^2^, C^3^), nor to the way BOLD or LFP was derived from the population activity – the simulation algorithm and the analysis code were identical for all simulations. The difference arises only from the different parameters – that is, there were different mixtures of C^1^, C^2^, and C^3^ for the V1 and the V2 simulations. This highlights the fact that the identical mechanism converting neural activity to BOLD (“neurovascular coupling”), modeled here as power in the time series summed across neurons (Equation 3), can produce very different correlations between BOLD and features of the LFP, depending on the neural activity.

#### 2.4.2 The relation between the BOLD amplitude and LFP features across sites

In the example V1 site and the corresponding simulation, the BOLD signal was well predicted by broadband increases (Fig 8B, Fig 8D). In the example V2 site and the corresponding simulation, the BOLD signal was predicted by both broadband increases and alpha decreases (Fig 8C, Fig 8E). By explicitly modeling both the population response and the population-to-instrument transformations, we see that a difference in the relation between instrument measures (BOLD and ECoG) can arise from a difference in the population response, without a difference in neurovascular coupling. We now ask (1) whether these effects are consistent across the measured V1 and V2/V3 sites, and (2) how a multiple regression model using broadband, gamma and alpha as predictors fits the BOLD response for both data and simulation.

As we argued in the Introduction, we believe there is no single, general transfer function that can predict the BOLD signal from the LFP. Yet a regression model linking the two measures can be a useful way to summarize the results of a particular experiment or simulation, and to compare results between different experiments or simulations. Here, we fit several regression models to each simulation and to the data (Fig 9). The regression models predicted the simulated or measured BOLD response from either a single LFP component (broadband power, gamma power, or alpha power), or from combinations of LFP components (each of the pairwise combinations, and the 3 components together). These regression models were fit separately for each of the 9 sites in V1 and each of the 13 sites in V2/V3, and for the 22 corresponding simulations. Accuracy of each model was assessed by the variance explained in the cross-validated data (*R*^*2*^, the coefficient of determination). With a cross-validation procedure, there is no advantage in accuracy for models with more free parameters, and accuracy is reduced rather than increased from overfitting. The cross-validated *R^2^* was compared to the *R*^*2*^ in a null distribution, derived from shuffling the assignment between data and stimulus conditions.

**Fig 9.**
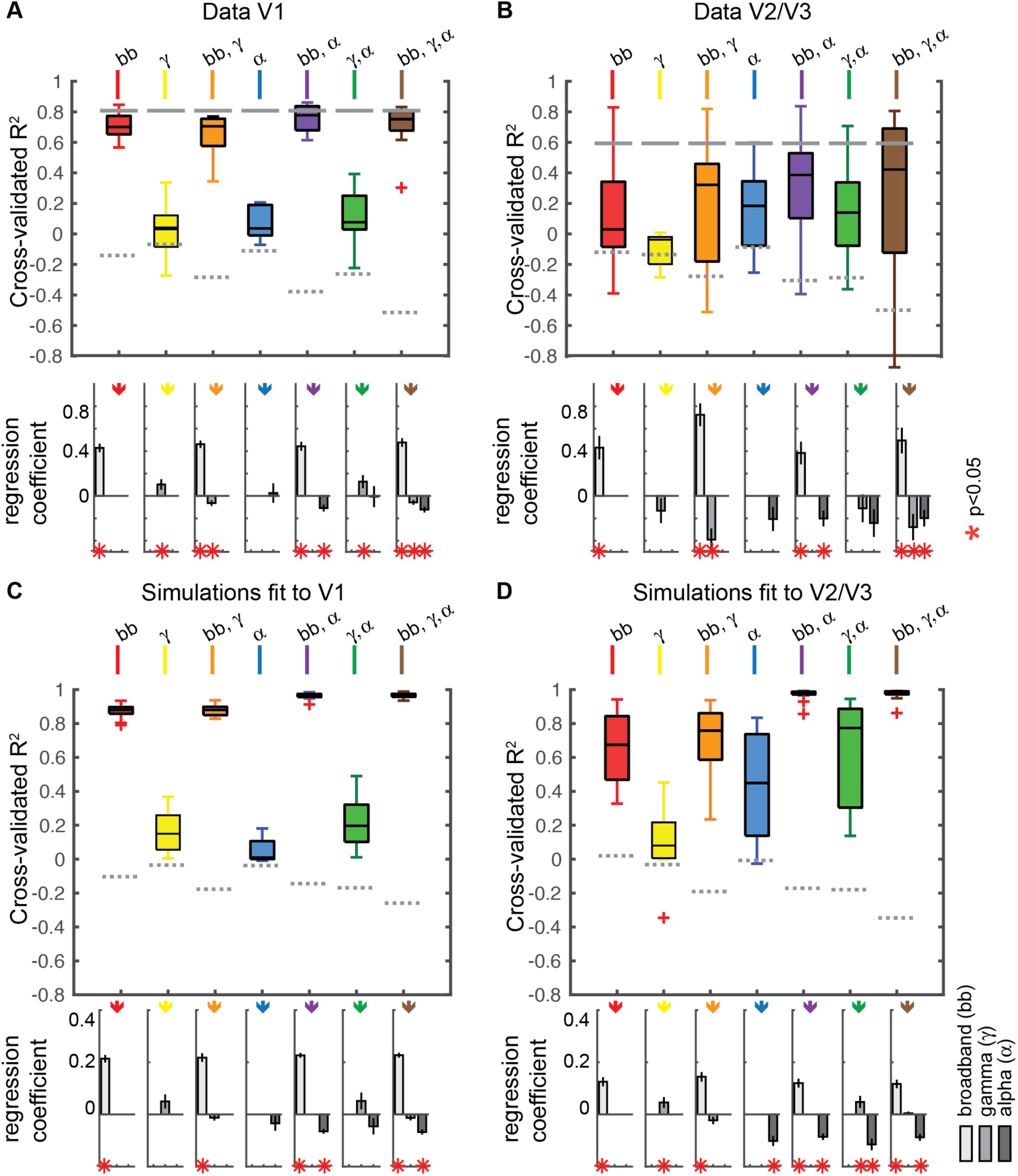
Explained variance in the BOLD fMRI signal in the simulations and in data. (A) Variance in the measured BOLD signal explained by broadband, gamma and alpha changes in the ECoG data. The colored box plots show the variance explained by each of the 7 model types: black bar = median, upper and lower boxes = quartiles, and error bars = data range excluding outliers (outliers plotted as red pluses). The *R*^*2*^ was cross-validated (split between subjects for BOLD and stimulus repetitions for ECoG), to ensure that the *R*^*2*^ can be compared between models with different numbers of explanatory variables. Gray dashed lines indicate the noise floor (*R*^*2*^ when shuffling the BOLD labels in the training data), and the solid gray lines indicate the data-to-data reliability for the BOLD signal. Bottom: The regression coefficients show whether the broadband, gamma, and alpha signals were positive or negative predictors of the BOLD signal. A red * in the lower plot indicates whether regression coefficients differed significantly from zero by a bootstrap test (p<0.05). (B) Same as A, but for the 13 V2/V3 electrodes. (C) Same as A, but for the 9 simulations fitted to V1 data. The *R*^*2*^ was cross-validated (split between even and odd stimulus repetitions). D) Same as C, except for the 13 simulations fitted to the V2/V3 data.

##### Data

The single parameter regression models across V1 electrodes (Fig 9A) show the same pattern as the single electrode examples in Fig 8: broadband alone was a good predictor of BOLD (median cross validated *R*^*2*^=0.70 across 9 sites) while gamma and alpha alone were not (gamma *R*^*2*^ = 0.04, alpha *R*^*2*^ = 0.04). Across all regression models, the observed BOLD signal was best predicted by a combination of broadband and alpha changes with an *R*^*2*^ = 0.78, close to the data-to-data reliability (*R*^*2*^ = 0.82). Adding gamma as a predictor to either the broadband only model (model 2 versus model 1) or to the combined broadband and alpha model (model 7 versus model 5) did not increase model accuracy, confirming that the gamma amplitude was not predictive of the BOLD signal.

The single parameter regression models across V2/V3 sites (Fig 9B) show that alpha power predicted the BOLD signal (*R*^*2*^ = 0.32, with a negative Beta value), whereas broadband alone was only slightly more accurate than a control, shuffled model, and gamma alone had no predictive power. As in V1, the BOLD response was best explained by a regression model combining broadband and alpha (*R*^*2*^ = 0.39; see also Fig S6), or a model using all three predictors (*R*^*2*^ = 0.42). Overall, compared to V1, the BOLD signal in V2/V3 was less accurately predicted by the regression models based on the electrophysiological measurements. As with the case of predicting BOLD from simulated neuronal activity, predicting BOLD from ECoG measures is limited by the reliability of the stimulus-evoked BOLD signal, which was lower for V2/V3 than for V1 (0.59 versus 0.82% *R*^*2*^).

##### Simulations

For the simulations, we expect broadband power to positively predict the BOLD signal and alpha power to negatively predict the BOLD signal, because of the construction of the simulations: broadband and alpha power elevations were achieved by increasing the variance per neuron, rather than correlations between neurons; the converse was true for gamma. Nonetheless, solving the regression models can be informative because, as seen in Fig 8, simulations with the identical input types and the identical analysis can lead to different patterns, depending on the parameters (weights) in the simulations. Moreover, the regression analyses of the simulated data can be compared against the regression analyses of the measured data.

The results from the regression model on simulated V1 LFP and BOLD (Fig 9C) were qualitatively similar to the V1 data: broadband alone was a good predictor of BOLD (median cross validated *R*^*2*^=0.70 across 9 simulations) while gamma and alpha alone were not (median *R*^*2*^ = 0.04 for both predictors). For simulations fit to V2/V3 ECoG data (Fig 9D), alpha alone predicted the BOLD signal with moderate accuracy (median *R*^*2*^ = 0.32).

Similar to the data, the BOLD response in the simulations fit to V1 and V2/V3 was well explained by a regression model combining broadband and alpha (*R*^*2*^ = 0.78, *R*^*2*^ = 0.39, Fig 9C **and** D. The regression coefficients for these models were positive for broadband and negative for alpha. A model that incorporated all three LFP measures – broadband, alpha and gamma – explained little to no additional variance for either simulation, confirming our earlier observation that narrowband gamma power was not correlated with BOLD amplitude in simulated data. The generally higher variance explained in V1 than in V2/V3 again matches the higher BOLD reliability in V1 for these experiments.

Across simulations and data sets, a general pattern emerges. The broadband signal was positively predictive of BOLD, and alpha power was negatively predictive of BOLD. Narrowband gamma had no consistent relation with BOLD. While the relationships between broadband and BOLD and between alpha power and BOLD were consistent in terms of sign (the former positive, the latter negative), the level was not always the same. As we noted in the example sites shown in Fig 8, and the summary across sites in Fig 9, the broadband power was more strongly predictive of BOLD in V1, and alpha power was more strongly predictive in V2/V3. An examination of responses to the different stimulus types clarifies the difference between V1 and V2/V3 in these data. Specifically, the BOLD response in V2/V3 to noise patterns was under-predicted by the broadband response alone (Fig S5). In V2/V3 alpha decreased more for the noise patterns and this alpha decrease accounted for the BOLD change unexplained by broadband (Fig S1). This helps to explain why a model that includes broadband and alpha is much more accurate for V2/V3 than a model that includes only broadband. In contrast, for V1 the BOLD response was well predicted by broadband power in most sites, with little systematic prediction failures, and little room for increased model accuracy when adding predictors such as alpha power.

#### 2.4.3 Correlation between BOLD and LFP across all frequencies

In the previous section, we modeled the BOLD responses as a linear function of three components derived from the LFP. These features – broadband power, narrowband gamma power, and narrowband alpha power – are summary metrics of the power spectrum. We also tested how the power at each frequency in the ECoG data and in the simulated LFP correlated with the BOLD response (measured and simulated, Fig 10).

**Fig 10.**
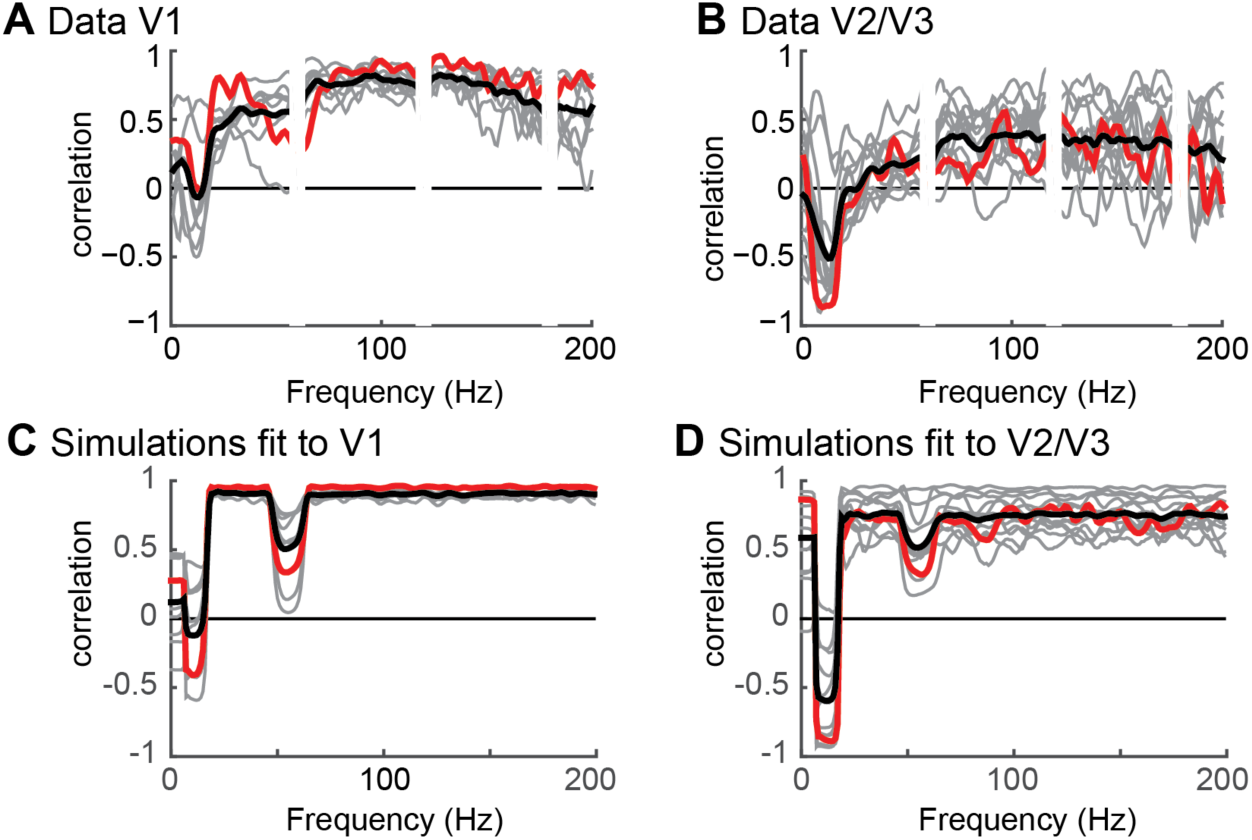
The correlation between BOLD and LFP as a function of frequency. (A) The correlation between ECoG and BOLD for the V1 data shows a positive correlation between ECoG and BOLD for a broad range of frequencies, except those including the alpha changes. Gray lines represent the 9 individual V1 electrodes, the black line is the average, the red line corresponds to the example sites shown also in Fig 7. (B) In the V2/V3 data there was a strong negative correlation between ECoG and BOLD in the alpha range around 10 Hz and a positive correlation between ECoG and BOLD for a broad range of frequencies. Gray lines represent the 13 individual V2/V3 electrodes, the black line is the average, the red line corresponds to the example electrode shown also in Fig 7. Note that neither the V1 electrodes nor the V2/V3 electrodes show a peak at the gamma frequency. (C) The correlation between LFP and BOLD for simulations fit to V1 shows that there is a positive correlation across most frequencies, except those including the alpha and gamma changes. (D) The correlation between LFP and BOLD for the simulations fit to V2/V3 shows that there is a strong negative correlation around 10 Hz, and a positive correlation across a broad range of frequencies.

##### Data

We calculated ECoG power for each frequency from 1 to 200 Hz and correlated the power changes from baseline with BOLD changes from baseline. In V1, ECoG responses across all frequencies except the alpha band were positively correlated with the BOLD response (Fig 10), consistent with the regression analyses of the summary metrics, showing that broadband ECoG power was the best predictor of the BOLD signal.

The pattern of correlation between ECoG power and BOLD in V2/V3 was similar to that found in V1, although the overall level of correlation was lower (Fig 10). There were positive correlations between ECoG and BOLD extending across most frequencies that were weaker than in V1, and there was a negative correlation for most sites in the alpha band.

##### Simulation

The correlation between simulated BOLD and ECoG power in V1 was qualitatively similar to that found in the data. In simulations fit to V1 ECoG data, the LFP correlated well with the BOLD signal across all frequencies except those in the alpha band (8-15 Hz) and below, and those in the gamma band (40-60Hz).

In simulations fit to V2/V3 ECoG data, the pattern was similar, except that the correlation was negative in the alpha band rather than 0, and weaker but still positive in the rest of the spectrum. These patterns match the summary metrics of alpha, gamma, and broadband shown in Fig 9.

##### Site to site differences

There were some differences between V1 sites. For example, in two sites, the correlation across frequencies dipped in the gamma band (30-80Hz), similar to simulated data. These are also the two sites that showed the largest amplitude gamma responses (sites 8 and 9 in Fig S5). In other words, when cortical sites showed large gamma signals, these signals were uncorrelated with BOLD. The fact that in 7 V1 sites there was a positive correlation between BOLD and LFP power spanning 30-80 Hz might seem inconsistent with our earlier observation that narrowband gamma power was not predictive of the BOLD signal in V1 sites (Fig 8 **and** Fig 9). However, in this analysis the narrowband and broadband power are not modeled separately and the positive correlation between power at 30-80 Hz in Fig 10 thus likely suggests that broadband power extends into this band, since broadband changes can extend across all frequencies [11,37]. Therefore, if there is little to no narrowband response, we would expect a positive correlation between BOLD and ECoG throughout all frequencies.

There were some site-to-site differences in the correlation between alpha and BOLD. For example, some sites showed a positive correlation with BOLD in the alpha range, and others showed a negative correlation (Fig 10A **and** B). These site to site differences depend on the range of responses evoked by stimuli. For example, for electrodes in which stimuli evoked a large of power changes in the alpha band, alpha was more strongly correlated with BOLD. Similarly, for electrodes in which stimuli evoked a large range of broadband responses, broadband was more highly correlated with BOLD (Fig S8). This pattern did not hold for narrowband gamma power changes.

## 3. Discussion

This study investigated the relationship between electrophysiological and BOLD measurements in human visual cortex. Our modeling framework decomposed the signals into two stages, a first stage in which we simulated the neuronal population responses (dendritic time series), and a second stage in which we modeled the transfer of the neuronal time series to the BOLD signal and the field potential. This approach differs from the direct comparison of electrophysiological signals and BOLD. The explicit separation into stages clarified both a similarity and a difference between the BOLD amplitude and the field potential power: the two can be approximated as the same operations on the neuronal population activity, but applied in a different order. Specifically, within a brief window, we modeled the BOLD amplitude as the sum of the power in the neuronal time series, and the field potential as the power of the sum of the neuronal time series. Because the order of operations differs, the two signals differ, and each is blind to particular distinctions in the neuronal activity. For example, the BOLD signal (according to our model) does not distinguish between synchronous and asynchronous neural signals with the same total level of activity. In contrast, the field potential does not distinguish counterphase responses from no response. Even if one knew the exact mechanism of neurovascular coupling and the precise antenna function of an electrode, one still could not predict the relationship between the BOLD signal and the field potential without specifying the neuronal population activity that caused both. Hence the relationship between the two types of signals is not fixed, but rather depends on the structure of the underlying responses of the neuronal population.

Although we do not have access to the complete set of individual neuronal responses in any of our experiments in visual cortex, we approximated the responses by specifying the type of signals common to visual cortex. We therefore limited the space of neuronal population responses by modeling the activity as arising from three types of signals, enabling us to compute the complete set of field potentials and BOLD responses to a variety of conditions. Finally, we compared the simulated patterns of BOLD and field potential responses to the actual responses we observed in data from human subjects. These patterns are discussed and interpreted below.

### 3.1 Changes in broadband power predict BOLD

Many studies have reported correlations between BOLD and power in the gamma band LFP (30-130 Hz) (review for human studies: [49]). Yet changes in gamma band power do not reflect a single biological mechanism. For example, several recent studies have emphasized that LFP power changes in the gamma band reflect multiple distinct neural sources, including narrowband oscillations and broadband power shifts, with very different stimulus selectivity and biological origins [35,50,51].

Broadband changes have been proposed to reflect, approximately, the total level of Poisson distributed spiking (or spike arrivals) in a local patch of cortex [37]. In contrast, the narrowband gamma response is caused by neural activity with a high level of cell-to-cell synchrony [52] and likely depends on specialized circuitry [53]. While the two responses are sometimes distinguished as ‘high gamma’ (referring to broadband signals), and ‘low gamma’ (referring to oscillatory signals), this distinction is not general. Broadband signals can extend into low frequencies [11,54] so that the two signals can overlap in frequency bands. Hence separating gamma band field potentials into an oscillatory component and a broadband (non-oscillatory) component is not reliably accomplished by binning the signals into two temporal frequency bands, one low and one high, but rather requires a model-based analysis, such as fitting the spectrum as the sum of a baseline power law (to capture the broadband component) and a log-Gaussian bump (to capture the oscillatory component) [35].

There is strong experimental support for the idea that increases in broadband LFP power primarily reflect increases in asynchronous neural activity rather than increases in coherence. First, experiments have shown that broadband power is correlated with multiunit spiking activity [51,55]. Second, unlike the case of narrowband gamma LFP, changes in broadband LFP are not accompanied by changes in broadband spike-field coupling ([40], their Fig 1A-B). The possibility that neuronal synchrony sometimes affects broadband signals cannot be ruled out, for example as shown in cases of pharmacological manipulations in nonhuman primate [56]. In such cases, there would not be a simple relationship between broadband power and BOLD.

The prior literature has not shown definitively whether broadband LFP, narrowband gamma, or both predict the BOLD signal. The first study that directly compared simultaneously recorded BOLD and electrophysiology showed that both LFP power in the gamma frequency range (40-130 Hz) and multi unit spiking activity (MUA) predicted the BOLD signal [16], and further, that when the LFP power diverged from MUA, the gamma band LFP predicted the BOLD signal more accurately than did spiking. This study however did not separately test whether a narrowband (oscillatory) or a broadband (non-oscillatory) component of the LFP better predicted the BOLD response.

Other studies, too, have shown a variety of patterns when correlating LFP power changes in the gamma band with BOLD. Some reported that BOLD amplitude correlates with narrowband gamma activity [13], while others showed that BOLD correlates with broadband changes [11], and many did not distinguish narrowband from broadband power in the gamma band [57]. Simultaneous recordings of hemodynamic and neuronal activity in macaque V1 showed that BOLD signals from intrinsic optical images can occur in the absence of gamma band LFP changes [58], and that in some circumstances, multiunit activity predicts the BOLD signal more accurately than gamma band LFP [24,59].

Here we separately quantified the broadband power (spanning at least 50-150 Hz) and narrowband gamma power. We found that the amplitude of broadband changes accurately predicted the BOLD signal in V1. The empirical results and the models help resolve the question of why ‘high gamma’ has been shown to correlate with BOLD, and ‘low gamma’ sometimes does not [24]. The likely reason is unrelated to the difference in frequency range, nor to the size of the spectral perturbation in the local field potential. In fact, the elevation broadband power is relatively small (2 or 3 fold) compared to the elevation in power often observed in narrowband gamma oscillations (10 x or more)[35]. Instead “High gamma” is predictive of the BOLD signal in many cases not because of the specific frequency range, but because this signal captures the level of asynchronous neuronal response; this signal happens to be most clearly visible in the high frequency range (>100 Hz) where it is not masked by rhythmic lower frequency responses. Hence the distinction in predicting the BOLD response is not about “high” versus “low” gamma, but rather synchronous versus asynchronous responses, and the broadband signal, sometimes labeled high gamma, maps onto the first term on the right hand side of Equation 4, the portion of the field potential measurement which sums the energy demand of each neuron.

Our model fits and data support this view. When we captured the stimulus-related broadband response by simulating a change in broadband coherence across neurons rather than a change in the level of response in each neuron, our predicted BOLD response was highly inaccurate (Fig S7).

### 3.2 Changes in narrowband gamma power do not predict BOLD

In contrast, we propose that ‘low gamma’ often does not predict the BOLD response because ‘low gamma’ reflects narrowband oscillatory processes, which largely arise from a change in neuronal synchrony across the population rather than a change in the response level per neuron. This corresponds to the second term in the right hand side of Equation 4, the portion of the field potential measurement which reflects the cross-power arising from currents in different neurons, and which in our model is independent of the signals giving rise to the BOLD signal.

Our results and model do not argue that narrowband gamma oscillations will never be predictive of the BOLD signal. If in a particular experiment narrowband gamma oscillations were to co-vary with broadband increases, we would expect both signals to correlate with BOLD. This might occur in an experiment with gratings of different contrast; with increasing contrast narrowband gamma responses, broadband responses, and BOLD responses all increase [21,60] and all 3 measures would correlate across stimuli. In such an experiment, if narrowband gamma oscillations had a higher signal to noise ratio than the broadband response, then the oscillatory signal would likely show a higher correlation with BOLD. In contrast, when the choice of stimulus or task can independently modulate broadband power and gamma oscillations so that the two LFP measures are not correlated, as in the experiments presented here and previously [35], then gamma oscillations will not strongly correlate with BOLD.

Our simulation and empirical results are consistent with studies which varied chromatic contrast and spatial frequency, while measuring MEG and BOLD. These studies found that BOLD and narrowband gamma activity did not match in stimulus specificity [18,19]. It is likely that these stimulus manipulations, like ours, independently modulated narrowband gamma power and broadband power, although the studies did not quantify broadband fields, which are more challenging to measure with MEG than with ECoG [61]. We speculate that broadband fields spanning the gamma range would have shown a higher correlation with BOLD.

### 3.3 Neuronal synchrony and the BOLD signal

In our model, the LFP measures are highly sensitive to neuronal synchrony, whereas BOLD is not. In our simulations, increases in neuronal synchrony drove narrowband gamma oscillations in the field potential. There are other cases of population activity with a high degree of neuronal synchrony. One example is the steady state evoked potential associated with a periodic stimulus [first reported by 62,reviewed by 63]. Previous studies have described discrepancies between evoked potentials and the BOLD signal, such as in the case of spatial summation [11], directional motion selectivity [7,8] and spatial attention [9,10]. Our modeling framework suggests that the neural sources generating the steady state potential (synchronous neural activity) are likely to be only weakly related to the BOLD signal (depending largely on asynchronous signals), as these sources will primarily affect the second term on the right hand side of Equation 4 (cross-power). This does not imply that the two measures are always or even usually discrepant; the BOLD signal and steady state potentials are likely to correlate any time that the steady state signals correlate with other measures of neural activity. When measures do dissociate, we do not conclude that one measure is more accurate; instead, the measures offer complementary views of the population activity, emphasizing the degree of synchrony or the average level of the response. An intriguing question is how each of the two signals contributes to perception and behavior.

Neural synchrony can also emerge without being time-locked to the stimulus, often called ‘induced synchrony’ or ‘induced oscillations’ [64]. In our simulation, we assumed that narrowband gamma LFP changes were induced by increases in *synchrony* between neurons, and not by changes in the *level* of gamma power within the individual neurons. In contrast, we assumed that broadband LFP increases were induced by increased broadband activity in individual neurons, and not by increased broadband coherence between neurons. (In Equation 4, a change in the left hand side, LFP power in the gamma band, can be produced by a change in either the first or second term on the right). This explains why, in our simulation, the broadband power was correlated with BOLD whereas the LFP gamma power was not, findings that were also confirmed by the data. Were our assumptions justified?

In principle, an increase in narrowband gamma power in the LFP could arise because the neurons synchronize in the gamma band, or because ongoing gamma oscillations within each neuron increase in amplitude, independent of coordination between neurons. There is strong experimental support for the former. Experiments which measure both intracellular membrane potential from single neurons and the extracellular LFP show that when there is an increase in narrowband LFP gamma power, the gamma power from individual neurons becomes more coherent with the LFP [44]. Moreover, the coherence between local spiking and the LFP also increases in the gamma band when LFP gamma power increases [40]. These results are consistent with our assumption that a significant part of the increase in gamma LFP power arises from a change in population coherence. To our knowledge, it is not certain whether there is also some increase in the level of gamma signals within individual neurons when the narrowband gamma band LFP power changes. However, since we can attribute a large part of the change in gamma LFP to a change in coherence, we infer that we can only attribute, at most, a small part of the change in gamma LFP to the level of gamma power within neurons.

In our simulation, we made two simple but extreme assumptions. First, we assumed that gamma oscillations occur with no change in the total level of neural activity, and hence no change in metabolic demand or BOLD. Second, we assumed that broadband responses occur with no change in neural synchrony. While these assumptions are likely incorrect at the limit, the simulations nonetheless captured the pattern of ECoG and fMRI results obtained in our datasets. Alternative models in which the broadband response was caused by a change in synchrony were much less accurate (Fig S7). Models in which gamma responses were caused by a change in level were only slightly less accurate, and cannot be ruled out entirely (Fig S7). However the regression models fit to our data (Fig 9) show that the power of narrowband gamma oscillations does not predict the BOLD response. Hence the most parsimonious explanation is that these responses in the LFP are caused in large part by changes in synchrony.

### 3.4 DC offsets and the BOLD signal

Both our measurements and our simulations showed that broadband electrophysiological responses were related to, but did not fully account for, the BOLD signal. This was especially evident in Simulation 2 and extrastriate data (V2/V3). In these cases, the amplitude of low frequency oscillations (8-15 Hz) was negatively correlated with the BOLD signal, independent of broadband signals. Numerous previous studies have reported that low frequency oscillations are anti-correlated with BOLD, including measurements in motor, visual and language areas [20-22,65-67]. This result may appear to conflict with the prior discussion, since we argued that oscillations (to the degree that they reflect neuronal synchrony) should have little to no effect on metabolic demand or the BOLD signal. It is therefore important to ask why low frequency oscillations sometimes correlate with the BOLD signal, both in data and in simulation.

One explanation is that alpha oscillations, or a mechanism which generates the oscillations, affect the BOLD signal indirectly, by inhibiting cortical activity. According to this explanation, an increase in alpha power results in a decrease in local spiking activity in turn reducing metabolic demand and the BOLD signal [68]. Alpha oscillations may indeed co-occur with reduced cortical excitation [69]. However, if this coupling between alpha power and spiking were the only explanation for the relationship between alpha power and BOLD, then a more direct measure of neuronal excitation, such as broadband or multiunit activity, would adequately predict the BOLD signal; alpha power would negatively correlate with the BOLD signal, but would provide no *additional* predictive power. Our data and model do not support this explanation, as we find that for most cortical sites, the most accurate predictor of the BOLD signal is a combined model including both the amplitude of alpha oscillations and broadband power.

We therefore propose that in addition to the indirect effect of modulating cortical excitability, alpha oscillations are also accompanied by a DC shift in membrane potential, making it less depolarized (i.e., closer to the equilibrium potential), and this shift reduces metabolic demand. Indirect evidence for a DC shift comes from MEG and ECoG studies [46,47,70], which refer to alpha oscillations as being asymmetrical (i.e., they are not centered at 0 – there is a DC shift). This can be explained by a simple process: if alpha oscillations reflect periodic inhibitory pulses, then on average they will cause a hyperpolarization (or less depolarization). If the neuron was slightly depolarized before the inhibitory alpha pulses, then the pulses would push the neuron toward equilibrium, and hence a lower energy state. In this view, large alpha oscillations reflect larger inhibitory pulses, reducing depolarization. We suggest that this reduced depolarization affects metabolic demand in two ways: by reducing spiking (as discussed above), and by maintaining a less depolarized state, reducing metabolic demand. In our model, the contribution to the BOLD signal from each neuron is the power in the time series (Equation 3), and the mean contributes to power. The idea that a DC shift in the membrane potential affects metabolic demand (in addition to altering excitability) is consistent with the observation that slowly changing currents (<0.5 Hz) correlate with BOLD fluctuations [12,71]. Moreover, if alpha oscillations are associated with a DC shift in membrane potential, this would explain why cortical excitability depends on the phase of the alpha cycle: at one phase the membrane potential is more depolarized, and hence cortex is more excitable, and in the opposite phase cortex is more hyperpolarized, and hence less excitable. This is consistent with the observations that the threshold for eliciting a phosphene with TMS changes with alpha phase [72,73] and that the alpha phase at the time of stimulus presentation influences the size of the BOLD response in visual cortex [74].

Inhibition takes two neurons – one to inhibit and one to be inhibited. In our simulations, the alpha oscillations (*C*^*3*^) were associated with inhibitory fluctuations in the membrane potential (mean below 0), which in turn was associated with decreases in BOLD. It is important to note that these fluctuations are meant to capture the effect of local inhibition on the post-synaptic neurons (the neurons being inhibited). The inhibitory neurons themselves are pre-synaptic, and the action of inhibiting other neurons is presumably an active process that consumes energy. Therefore inhibition is expected to increase energy demand in some neurons (the pre-synaptic neurons) and decrease energy demand in other neurons (post-synaptic neurons). We did not model the inhibitory neurons explicitly; however, the neural activity associated with active inhibition would be expected to contribute to the measured broadband signal in the ECoG data, and is implicitly included in the broadband inputs in our simulations (*C*^*1*^). More complex models (see paragraph 3.6) in which the circuitry of excitatory and inhibitory neurons is explicitly represented (such as [60,75,76]), may provide insight into how the balance between excitation and inhibition influences the field potential and the BOLD signal.

### 3.5 A single modeling framework accounts for patterns of LFP/BOLD correlations across sites

We found that the relationship between the BOLD signal and features of the ECoG data differed across cortical areas. For example, broadband changes in ECoG responses explained more variance in the BOLD data in V1 than in V2/V3. Conversely, low frequency power decreases (alpha, 8-13Hz) explained more variance in the BOLD signal in V2/V3 than in V1. In the absence of a model, we might have interpreted the results as evidence that neurovascular coupling differs across sites. Many previous studies have reported differences in the relation between LFP and BOLD as a function of site or condition, for example between cortical and subcortical locations [77], across cortical regions [78,79], between cortical layers [80], and as a function of medication [81]. Here, we showed that a difference in the relationship between LFP and BOLD need not arise because of a difference in neurovascular coupling. In our results, Simulations 1 and 2, like V1 compared to extrastriate areas, showed differences in the relationship between LFP and BOLD, yet we used the identical model of neurovascular coupling in all simulations. The systematic differences in the two simulations arose because of differences in the neuronal population activity, not because of differences in neurovascular coupling. While our results do not exclude the possibility of differences in neurovascular coupling across locations or states they do caution against interpreting differences in the relationship between field potentials and BOLD as evidence for a difference in neurovascular coupling, since they show that a single model can account for a variety of patterns. More generally, the V1 versus V2/V3 discrepancies bolster the argument that one cannot predict the exact relationship between BOLD and field potentials without also specifying the neuronal population activity.

### 3.6 The role of a simple model in understanding the relation between BOLD and LFP

A complete description of the biophysical processes giving rise to the BOLD signal and the field potential is far beyond the scope of this paper, and is likely premature given the enormous complexity in the nervous system, the vascular system, and the coupling mechanisms between them. Instead, the purpose of our modeling framework was to first begin with a general principle, namely that BOLD and field potentials sum neural activity according to a different sequence of operations; second, to instantiate this principle in simple mathematical rules; third, to combine these rules with a minimal model of neural population activity; and finally, to ask to what extent such a model can account for the patterns in our data. Our model omits many biophysical components, such as compartmentalized neurons, multiple cell types and vessel types, neurotransmitters, the dynamics of blood flow, and so on, and hence it is not a detailed simulation of the nervous system or vascular system. On the other hand, the simplicity of the model facilitates an understanding derived from basic principles, similar to the advantages in building computational, rather than biophysical, models of neural responses [82-85]. Both types of models and empirical studies are valuable. Here we emphasize that even with a highly simplified model of the BOLD signal, the field potential, and neuronal population activity, we are able to reconcile a wide range of findings in a complicated and technical literature. The model accounts for differences in how broadband field potentials and gamma oscillations relate to the BOLD signal. It can explain differences between cortical areas in the relationship between field potentials and BOLD. The model also provides an explanation for why the amplitude of alpha rhythms is negatively correlated with BOLD, even after accounting for the relationship between broadband signals and BOLD. We note that drastic simplifications are the norm in many fields of neuroscience, such as receptive field modeling of visual neurons; most such models omit fixational eye movements, optical properties of the eye, retinal and cortical circuitry, etc., instead modeling responses as a few simple mathematical computations of the stimulus (filtering, thresholding, and normalization) [86]. These highly simplified models will certainly fail under some conditions [87], yet they have proven to be of immense value to the field [85], in part due to their simplicity, and in part because the alternative, in which the responses of visual neurons are computed from a complete, neurobiologically realistic model of the nervous system simply do not exist.

### 3.7 Reproducible computations

To test competing computational theories about the relation between the visual input, the LFP and the BOLD response, it is essential to make sample data and code available for others [35,50]. Following standard practices of reproducible research [88-90], the Matlab code of the simulation, and sample data and code to reproduce the Figs in this manuscript can be downloaded at https://github.com/WinawerLab/BOLD_LFP.

### 3.8 Conclusions

To understand how the electrophysiology and BOLD responses are related, it is necessary to specify both the manner in which population activity transfers to the two signals, and the neuronal population activity itself. The former shows that the covariance between neuronal time series has a large influence on the field potential and not the BOLD signal. Based on our simulations and empirical results, we made several inferences about the neuronal population responses mediating the BOLD signal and the LFP: that narrowband gamma oscillations in visual cortex likely arise more from synchronization of neural responses than a change in level of the neural response, and hence have a large influence on the field potential and little influence on the BOLD signal; that responses which are asynchronous across neurons manifest in broadband field potentials and an elevated BOLD signal; and that low frequency oscillations observed in field potentials are likely accompanied by a widespread hyperpolarization, which in turn reduces metabolic demand and the BOLD signal. Our model-based approach brings us a step closer to a general solution to the question of how neural activity relates to the BOLD signal.

## 4. Materials and Methods

### 4.1 Simulated neuronal time series

Simulations were computed for a population of 200 neurons. Each simulation trial was 1 second long with millisecond sampling. The time series for each neuron was derived by summing three inputs, each 1 second long, followed by leaky integration with a time scale of 10 ms to simulate temporal integration in the dendrite (Fig 4). Each simulation was fit to ECoG data from one electrode and consisted of 240 trials, 8 repeats of 30 stimulus conditions. A condition in the simulation was defined by the parameter settings for the 3 inputs (Fig 4): C^1^ (broadband), C^2^ (gamma) and C^3^ (alpha). Variations in these three inputs resulted in power changes in the broadband, gamma, and alpha LFP. The inputs were fit to data such the simulated LFP power changes matched the ECoG data power changes for a particular electrode and stimulus.

#### 4.1.1 C^1^ - Broadband input

The first input was a series of random numbers drawn from a normal distribution, with no temporal dependencies and no dependencies between neurons.

*Motivation.* This input approximates spike arrivals with a Poisson distribution at a fixed rate for a given 1-s trial. A random normal distribution was used rather than a Poisson distribution for computational efficiency. (The pattern of results is the same for either distribution.) The input has a flat (white) power spectrum up to the sampling limit of 500 Hz. When coupled with leaky temporal integration (described in a subsequent section), this input results in a power spectrum that is approximately proportional to 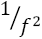 (brown noise). Several groups have proposed that the approximately 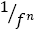 power spectra observed in field potentials arises from white noise (or Poisson noise) input to individual neurons, coupled to one or more low-pass filters [37,91,92]. Previously proposed sources of filters include an exponentially decaying current response in the synapse following each spike arrival [91], leaky temporal integration in the dendrite [37], and frequency dependent propagation in the extracellular tissue [93], the last of which has since been shown to be unlikely [94]. Regardless of the source of the low-pass filtering, the general proposal makes an interesting prediction, namely that a spectrally broadband increase in field potential power in response to a stimulus is likely to indicate an increase in the rate of spike arrivals following that stimulus [37]. This hypothesis has empirical support, based on correlations between spike rates (single unit and multiunit) and broadband field potentials [51,55], and the fact that a 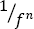 baseline spectrum, as well as stimulus-dependent broadband power increases, can be observed in intracellular (single neuron) membrane potentials in awake macaque visual cortex [39]. This hypothesis is the logic behind our choice to model both the baseline 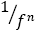 spectrum and stimulus-dependent broadband modulations as arising from spectrally flat inputs followed by low-pass filtering within individual neurons. For computational tractability, we explicitly modeled only one of the low pass filters – leaky integration in the dendrites. We assumed that spectrally broadband signals reflect uncorrelated activity. First, we have shown that the broadband ECoG signal is asynchronous with respect to a visual stimulus, and hence uncorrelated from trial to trial [11]. Here, we extrapolate that within a trial, the contribution to the broadband signal is asynchronous from neuron to neuron. One reason to assume so is based on a physiological model: the broadband signal has been hypothesized to arise from the leaky integration of Poisson distributed spike arrivals [37]. Even if the spike rate is correlated between neurons, the spike timing within a trial is likely have low correlations between neurons.

*Parameters.* For each simulation the Gaussian distribution defining C^1^ always had a mean, μ =.0.25. The slightly positive mean ensured that in the baseline state, the membrane potential was slightly positive, such that a suppressive signal (described in the section *C*^*3*^) could bring the potential closer to 0, hence reducing the metabolic demand. For the 8 conditions in each simulation, the baseline standard deviation of the distribution was set at σ = 0.3. A larger σ results in a larger broadband signal, and can be thought of as reflecting a higher Poisson rate of spike arrivals. The σ for each condition was calibrated such that the resulting changes in broadband power for each of the 8 stimulus conditions matched the changes in broadband power in the ECoG data (Fig 6, see Fig S9).

#### 4.1.2 C^2^ – Narrowband oscillations in the gamma band

The second input was band-passed filtered white noise. The white noise was drawn from a distribution with zero mean and fixed standard deviation on all trials and for all neurons, and subsequently band-pass filtered. Unlike C^1^, there were dependencies (coherence) between neurons. The level of coherence varied across the 8 trial types in each simulation.

*Motivation.* This input approximates a circuit producing narrowband gamma oscillations in the field potential. Parvalbumin positive interneurons project to pyramidal neurons, and can produce fluctuations in the membrane potential of the pyramidal neurons in gamma frequencies from 30-80 Hz [44]. The narrowband rise in gamma power associated with certain stimuli or tasks appears to reflect an increase in synchrony between neurons in this band [95]. Therefore, unlike C^1^, which varied in level but not coherence as a function of condition, C^2^ varied in coherence but not level as a function of condition.

*Parameters.* For all trials and all conditions, the white noise samples were drawn from a normal distribution with *μ* = 0 and *σ* = 0.*2*. The covariance of the distributions could range between 0 and 1 (using Matlab’s mvnrnd function). The white noise inputs were filtered between 50 Hz and 60 Hz prior to temporal integration in the dendrite: inputs were first zero-padded, then filtered with a 10^th^ order Butterworth filter in forward and reverse direction. (Fig 4). The covariance for each simulation was calibrated such that the resulting changes in narrowband gamma power for each of the 8 stimulus conditions matched the changes in narrowband gamma power in the ECoG data.

#### 4.1.3 C^3^ – Narrowband oscillations in the alpha band

The third input was band-passed filtered white noise, with an added asymmetry such that increased power decreased the mean amplitude. The coherence was the same for all trials and all neurons; the amplitude of the pulses varied by condition.

*Motivation.* Oscillations in the alpha band (8-15 Hz) are widely observed in visual cortex, with higher amplitudes associated with low sensory stimulation (e.g., eyes closed or zero contrast) or a low level attention. One model of alpha oscillations is that pyramidal neurons in visual cortex receive pulses of inhibition (hyperpolarizing inputs) spaced on the order of 100 ms, generated indirectly by thalamic-cortical loops [46,47]. According to this view, less active states are associated with larger inhibitory pulses, resulting in a time-averaged hyperpolarization, compared to more active states with smaller inhibitory pulses. The inhibition is pulsed rather than continuous, so that the reduced cortical responsiveness is dependent on the phase of the alpha cycle (most reduced following each inhibitory pulse). Individual neurons in visual cortex can indeed show membrane oscillations at frequencies around 10 Hz [48], indicating that it is reasonable to model the alpha oscillation measured in the population as arising from oscillations in individual neurons, rather than arising only from band-limited coherence between neurons.

*Parameters*. For all trials and all conditions, the white noise samples were drawn from a normal distribution with μ = 0 and σ = 1. The covariance of the distributions was fixed at.75 (using Matlab’s mvnrnd function). The white noise inputs were filtered between 9 Hz and 12 Hz prior to temporal integration in the dendrite: inputs were first zero-padded, then filtered with a 10^th^ order Butterworth filter in forward and reverse direction. The envelope was calculated by a Hilbert transform, and added to the signal, and the signal was multiplied by -1, such that increases in power reduced the mean amplitude. The signal was then multiplied by a factor c_3_, which was calibrated such that the resulting changes in narrowband alpha power for each of the 8 stimulus conditions matched the changes in narrowband alpha power in the ECoG data.

#### 4.1.4 Fitting the LFP power changes to the ECoG power changes

Changing inputs in C^1^, C^2^ and C^3^ results in a change in LFP power in broadband, gamma and alpha respectively. In order to fit the simulated LFP power changes to the ECoG power changes we quantified the input to LFP output relation, such that a certain change in simulated LFP power could be predicted by change in the input amplitude. Different functions described the relation between the input and LFP. The relation between broadband input and LFP broadband was described as

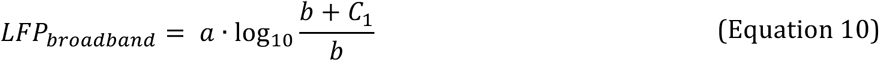

Since gamma and alpha were dependent on broadband amplitude (an increase in broadband noise masks the relative contribution of narrowband oscillations) the following function described the relation between input and gamma and alpha LFP:

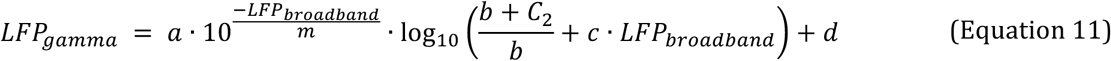

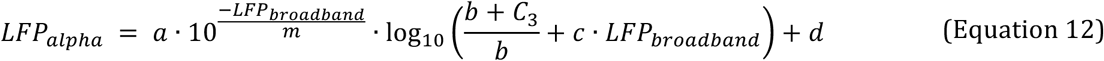

Parameters a, b, c, d and m were estimated by a separate calibration procedure in which C^1^, C^2^ and C^3^ were systematically varied and LFP broadband, gamma and alpha were calculated. Fig S9 shows that using this procedure the simulated LFP power changes match the ECoG power changes well.

### 4.2 Stimuli and task

Stimuli for ECoG experiments were reported previously [35]. In brief, for one subject, the stimuli came from 8 classes of patterns (30 exemplars per class, 20x20°), including high contrast vertical gratings (0.16, 0.33, 0.65, or 1.3 cycles per degree square wave) noise patterns (spectral power distributions of *k*/*f*^4^, *k*/*f*^2^, and *k*/*f*^0^), and a blank screen at mean luminance (Supplemental methods and Fig S2). For the second ECoG subject, there were the same 8 classes as well as two other stimulus classes – a high contrast white noise pattern and a plaid at 0.65 cpd. The fMRI subjects had the same 10 stimulus classes as the second ECoG subject.

#### 4.2.1 ECoG task

ECoG data were re-analyzed from a previous report [35]. We briefly summarize that experiment here. Subjects viewed static images of gratings and noise patterns for 500 ms each, with 500 ms of zero-contrast (mean luminance) between successive stimuli. Order of presentation was randomized (Fig S2). There were a total of 210 contrast stimuli, shown once each in a single, continuous experiment (and 210 interstimulus blanks). Stimuli were shown on a 15-inch MacBook Pro laptop using Psychtoolbox (http://psychtoolbox.org/). The laptop was placed 60 cm from the subject’s eyes at chest level. Screen resolution was 1280x800 pixels (33x21 cm). Coordinates of the population Receptive Fields (pRF) were obtained from a prior study [11].

#### 4.2.2 fMRI task

The fMRI experiment was a block design, with 12-second stimulus blocks alternating with 12-s blank periods (mean luminance). During the stimulus blocks, images were presented at the same rate as the ECoG experiment: 500 ms duration alternating with 500 ms of zero-contrast (mean luminance) between images (Fig S2). All stimuli from each block came from one of the 7 stimulus classes. The exemplars within the block were all different. Subjects participated in 8 scans of 9 blocks each, and block order was randomized using Latin squares. Two subjects (S2 and S3) additionally participated in an identical experiment using lower contrast images, resulting in similar findings. fMRI subjects participated in two pRF runs to identify retinotopy maps. Stimuli for the pRF experiments consisted of a bar (width= 3 deg) that swept across the visual field in 8 directions: the four cardinal directions and the four diagonals. The bar contained a drifting checkerboard with 100% contrast. Images were projected on a screen in the rear of the magnet bore using an LCD projector (LC-XG250, Eiki) with a resolution of 1024x768 (60 Hz refresh rate) and subtending approximately 32x24 visual degrees (32.4x24.3 cm). Subjects viewed the screen with a mirror mounted to the RF coil. The viewing distance was approximately 58 cm.

### 4.3 ECoG procedure

ECoG data were measured from two subjects who were implanted with subdural electrodes (2.3 mm diameter, AdTech Medical Instrument Corp) for clinical purposes at Stanford Hospital. Informed, written consent was obtained from all subjects. ECoG protocols were approved by the Stanford University IRB. In 22 electrodes in V1 V2 and V3, broadband and narrowband gamma responses were quantified as before [35], and alpha power changes were calculated (see Supplementary Methods).

#### 4.3.1 ECoG recording

ECoG data were recorded at 3052/1528 Hz (ECoG subject 1/ECoG subject 2) from 118/96 electrodes through a 128-channel Tucker Davis Technologies recording system (http://www.tdt.com). Electrodes were localized on a postoperative computer tomography (CT) scan that was co-registered with a pre-operative MRI, and locations were corrected for the brain shift [6]. Electrodes that showed large artifacts or epileptic activity (as determined by the patient’s neurologist) were excluded from analysis (7/35 electrodes were excluded in subject 1/subject 2). Off-line, data were re-referenced to the common average, low-pass filtered and resampled at 1000Hz for computational purposes using the Matlab resample function. Line noise was removed at 60, 120 and 180 Hz using a 3^rd^ order Butterworth filter.

#### 4.3.2 ECoG analyses

Broadband and narrowband gamma responses were quantified as before [35]. We calculated power spectra and separated ECoG responses into broadband and narrowband gamma band spectral power increases. To control for the influence of evoked activity on the spectrum, event related potentials (ERPs) were calculated per condition and the condition specific ERP was regressed from each trial. This procedure makes sure that the broadband increase is not due to a sharp edge in the ERP; the same pattern of results is obtained if this step is omitted. For each condition, the average power spectral density was calculated every 1 Hz by Welch’s method [96] from 0 – 500 ms after stimulus onset (and 0-500 ms after stimulus offset for the baseline) and a 250 ms Hann window to attenuate edge effects. ECoG power spectra are known to obey a power law and to capture broadband and narrowband gamma increases separately the following function was fitted to the average log spectrum from 35 to 200Hz (leaving out 60Hz line noise and harmonics) from each condition (Fig 3):

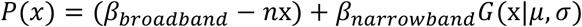

In which,

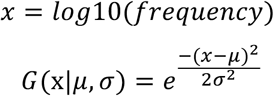

with 10^*σ*^ = 1.1 Hz and 35 Hz < 10^*μ*^ < 80 Hz.

The slope of the log-log spectral power function (*n*) was fixed for each electrode by fitting it based on the average power spectrum of the baseline. For cross-validation, trials were split into even and odd repeats, and broadband and gamma changes were calculated for each. Confidence intervals were calculated by a bootstrap procedure. For each condition C with N_c_ trials, N_c_ trials were drawn randomly with replacement and power spectra were averaged. The parameters β were fitted on the average log power spectrum from these bootstrapped trials. This was repeated 100 times, resulting in two sets of distributions of broadband and gamma weights for even and odd trials.

Alpha response amplitude was calculated as follows. Alpha changes are best visible after the initial onset transient and ERP, and we used the power from 250-500 ms to calculate the alpha decreases for each stimulus. Alpha amplitude was calculated by averaging the log-power between 8 and 13 Hz.

#### 4.3.3 Electrode selection

Electrodes for analysis were selected on the basis of three criteria. First, the pRF was located within V1, V2, and V3. Second, the explained variance in a pRF experiment was >15% [11]. Third, the center of the pRF was within the extent of the stimulus (<12 deg) and on the contralateral visual field. Because ECoG subject 2 did not have pRF data, only anatomical estimates of V1, V2, and V3 were used [97]. These criteria yielded 22 electrodes (19 from ECoG S1, 3 from ECoG S2).

### 4.4 fMRI procedure

Functional MRI data was measured from four subjects (3 female, ages 22-42) with normal or corrected-to-normal vision at the Center for Brain Imaging at NYU. Informed, written consent was obtained from all subjects. The fMRI protocols were approved by the New York University IRB. Functional MRI data were preprocessed and analyzed using custom software (http://vistalab.stanford.edu/software) (see Supplementary Methods). Disc ROIs (radius = 2 mm) were defined in fMRI subjects to match the position of the electrodes in ECoG subjects using a combination of anatomy, pRF centers, and visual field maps. The similarity between the ROI position and electrode position was compared via visual inspection of anatomical images and pRF centers (Fig S3).

#### 4.4.1 fMRI recording

Anatomical and functional MRI data was collected at the Center for Brain Imaging at NYU on a Siemens Allegra 3T head-only scanner with a Nova Medical transmit/receive coil (NMG11) and a Nova Medical phased array, 8-channel receive surface coil (NMSC072).

Two to three T1-weighted whole brain anatomical scans (MPRAGE sequence) were obtained for each subject (voxel size: 1x1x1 mm, TR: 2500 ms; TE: 3.93 ms, flip angle: 8 deg). Functional images were collected using gradient echo, echo-planar imaging (voxel size: 2x2x2 mm, 24 slices, TR: 1500 ms, TE: 30 ms, flip angle: 72 deg). Images were corrected for B0 field inhomogeneity during off-line image reconstruction using a separate field map measurement made half way through the scan session. Slice prescription was set approximately perpendicular to the calcarine sulcus, covering the occipital lobe. In addition, a T1-weighted inplane was collected with the same slice prescription to align functional images to the high-resolution anatomical images.

#### 4.4.2 fMRI analysis

*Preprocessing.* Anatomical images were co-registered and segmented into gray/white matter voxels using FreeSurfer autosegmentation algorithm (surfer.nmr.mgh.harvard.edu). A 3D mesh of the cortical surface was inflated for ease of visualization. Functional data were preprocessed and analyzed using custom software (http://vistalab.stanford.edu/software). Data were slice-time corrected to adjust for differences in acquisition time among slices in the 1.5-sec frame. Data were motion corrected for both between- and within-scan motion. Finally, data were high-pass filtered for low frequency drift [98] by multiple moving average smoothing (2 iterations, 40 seconds). Data were then converted to percent signal change by dividing each voxel’s signal by its mean signal. The first four frames of each run (6 sec) were discarded to allow longitudinal magnetization and the hemodynamic response to reach steady state.

*Analysis.* Noise was removed from the fMRI data using GLMdenoise, a variant of the standard GLM commonly used in fMRI analysis [99]. In brief, GLMdenoise derives noise regressors for each subject by performing principle components analysis on noise voxels that are unrelated to the task. The optimal number of noise regressors is selected based on improvement in cross-validated *R*^*2*^. The final model is fitted to each voxel’s time series and bootstrapped 100 times over 8 runs. Here, the predictors in the GLM were the nine image categories (4 gratings, 4 noise patterns, 1 plaid) and a blank period (a randomly assigned blank block). Voxel bootstraps were averaged across voxels within a region of interest (ROI). The resulting 100 bootstraps per ROI were vector-length normalized and averaged across subjects. The beta estimate for each condition is taken as the median averaged bootstrap and the standard error as one-half the 68% confidence interval.

*Population receptive field (pRF) model.* The pRF runs were analyzed by fitting a 2D Gaussian to each voxel, modeling its pRF [100]. The pRF is defined by center location (x,y coordinates) and spread (sigma). The resulting maps were used to define retinotopic areas V1, V2, and V3 as in [101].

### 4.5 Predicting fMRI signals directly from ECoG models

The relationship between fMRI and ECoG signals was analyzed using a linear regression model. The cross-validated coefficient of determination (*R*^*2*^) was used as a metric for model accuracy and the regression coefficients were used to test whether ECoG predictors (broadband, gamma and alpha) had a positive or negative relation with BOLD (also see Supplementary Methods).

The relationship between fMRI and ECoG signals was analyzed using a linear regression model:

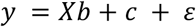

where *y* is a vector of fMRI amplitudes (beta estimates), with *n* entries for the *n* different stimuli; *X* is a matrix of ECoG responses, *n* by 1, 2 or 3, where the columns correspond to one or more of broadband, gamma and alpha estimates; *b* are the 1, 2 or 3 beta weights for the broadband, gamma, and alpha estimates; *c* is a constant (the *y-intercept*); and ε is the residual error term. The model was fitted separately for each cortical site (electrode/ROI pair), and for different combinations of predictors – broadband alone, gamma alone, alpha alone, each pairwise combination, and all three predictors together.

Models were evaluated by split-half cross-validation. First, the regression model *y*_*i*_ = *X*_*i*_*b*_*i*_ + *c*_*i*_ + *ε* was fit using half of the fMRI subjects (1 and 2) and half of the ECoG stimulus repetitions (even repetitions). To cross validate this model, the beta values (*b*_*i*_) were then applied to the left out half of the ECoG data (odd stimulus repetitions) to predict the left out half of the fMRI data (fMRI subjects 3 and 4). The same procedure was applied by reversing the training and testing data. This resulted in two testing data sets with BOLD responses predicted from ECoG for each stimulus condition (*X_i_b_i_* + *c_i_*) and an actual measured BOLD value. For each cortical site, the coefficient of determination (see below) was calculated between the concatenated predictions and BOLD data values of the two test sets. All *R*^*2*^ values reported in the results are cross-validated in this manner. The same pattern of results was achieved if instead of cross-validation, we solved the models on the complete data sets and computed the *R*^*2*^ adjusted for the number of regressors.

To test whether different ECoG predictors (broadband, narrowband, alpha) had a positive or negative relation with BOLD, we tested whether the regression coefficient was significantly larger or smaller than zero. The regression coefficient was considered to be significantly different from zero using a bootstrap statistic: for each model the median of the beta values across sites was calculated after resampling 10000 times. If <2.5% of the resampled statistics were smaller than zero, the beta values were considered significantly positive, and similarly, if <2.5% of the resampled statistics were greater than zero, the beta values were considered significantly negative.

### 4.6 Model accuracy

All model predictions were quantified using the coefficient of determination on cross-validated predictions. For predicting BOLD data from simulations of population neuronal activity (Fig 7, Fig S7), the predicted BOLD has arbitrary units. In these cases, the observed BOLD and the predicted BOLD were both normalized by subtracting the mean and then dividing by the vector length. When predicting BOLD responses from features of the LFP data (broadband, gamma, and alpha) by regression, the predicted BOLD data were in the same units as the measured BOLD, and no normalizing or re-scaling was done.

To quantify the accuracy of the models, we calculated the cross-validated coefficient of determination, *R*^*2*^:

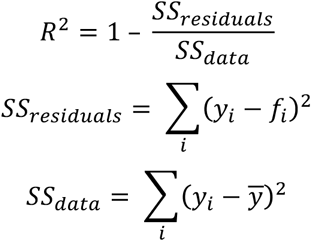

where *y* are the data values and *f* are the prediction values. Because the model fits are cross-validated, it is possible for the model errors (residuals) to be larger than the data values, hence *R*^*2*^ can be lower than 0, and spans -∞, 1. In the case in which the model predictions and the data are unrelated and each are normally distributed with equal variance, *R*^*2*^ will tend to –1.

## Acknowledgements

We thank two colleagues for making important contributions to our analytic framework. David J Heeger (NYU) proposed to us that the two measures, BOLD and LFP, could be approximated by the same operations computed in a different order (captured in Equations 1-3) and Weiji Ma (NYU) derived a variant of Equation 4 for us, expressing analytically the quantitative difference that results from the difference in the order of operations. This work was supported by the National Eye Institute at National Institutes of Health grant R00-EY022116 and the National Institute of Mental Health at National Institutes of Health grant R01MH111417-01 to J.W. and the National Eye Institute at National Institutes of Health grant T32-EY20485 and the Netherlands Organization for Scientific Research Veni grant 016.VENI.178.048 to D.H. The ECoG data, which was previously published and re-analyzed for this paper, was provided as part of the Stanford Human Intracranial Cognitive Epilepsy Program (SHiCEP), headed by Josef Parvizi. We also thank Brian A. Wandell, David J, Heeger, Rachel Denison, Noah Benson, and Nathan Witthoft for helpful comments on an earlier draft of the manuscript. The empirical results were presented at VSS in 2015 and the modeling components have been presented at SfN 2015.

## Supplemental materials to: Neuronal synchrony and the relation between the BOLD response and the local field potential

**Figure S1.**
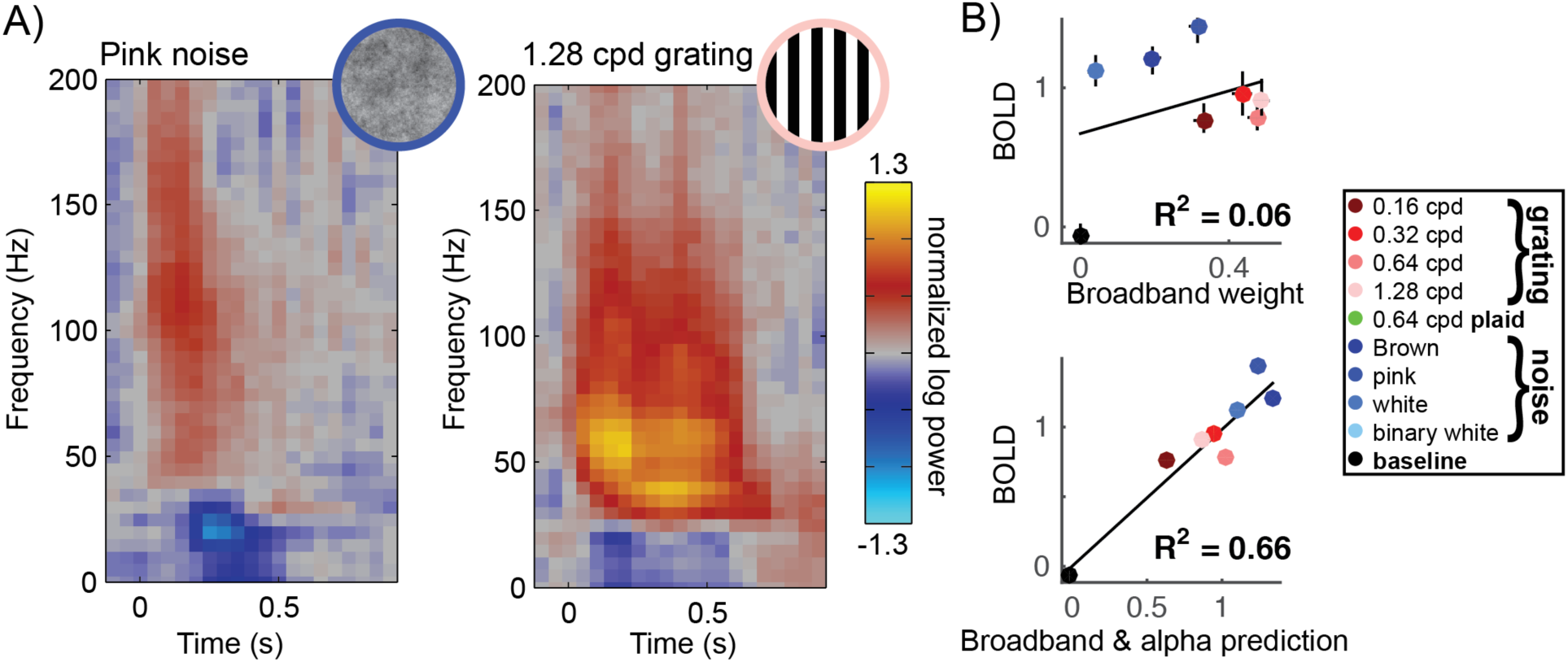
Alpha changes explain additional variance in the BOLD response. **A)** Time/frequency spectrograms for the pink noise pattern and the grating in an exemplary V2 electrode show that power in the low frequencies decreased more for the pink noise pattern (left) than for the grating (right). **B)** Top: the correlation between broadband and BOLD shows that the broadband response under-predicts the BOLD response for the noise patterns (blue dots). Red and pink dots represent the gratings. This pattern is visible in most V2/V3 electrodes (Figure S5) Bottom: taking into account the alpha decreases in the regression model explains the variance in the BOLD response that was not explained by the broadband changes. The R^2^ represents the cross-validated coefficient of determination.

**Figure S2.**
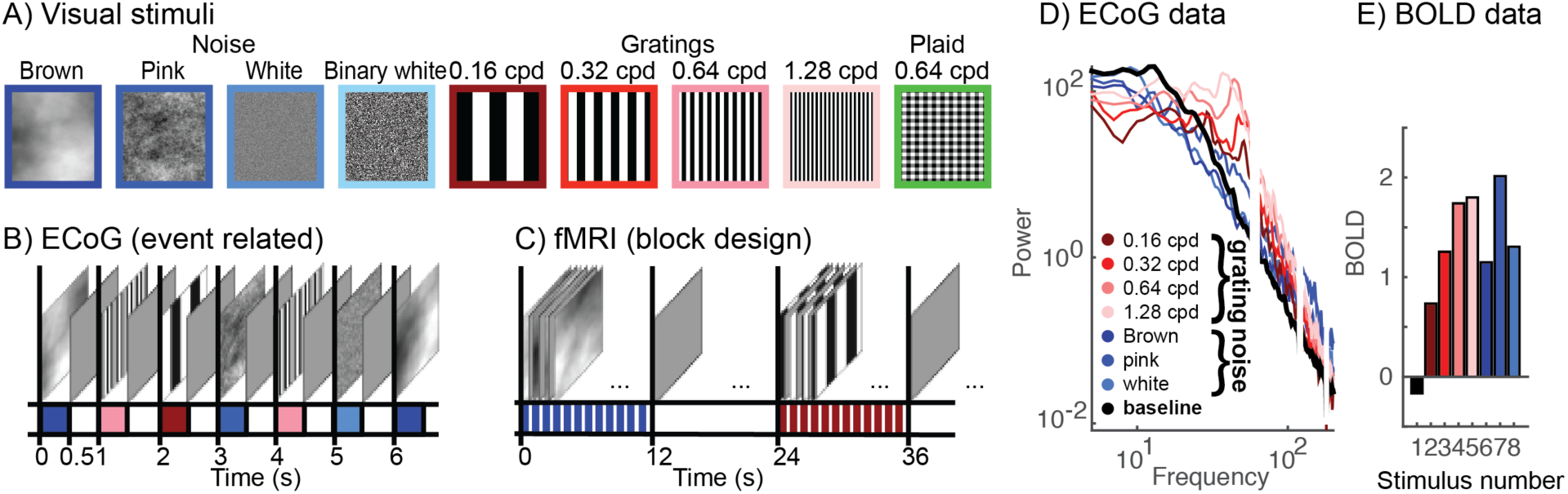
ECoG and fMRI measurements. **A)** ECoG and fMRI responses were measured to 8 different stationary stimuli. In all experiments, subjects were instructed to fixate on a dot at the center of the screen that alternated between red and green, changing colors at random times. Subjects pressed a button when the fixation dot changed color. ECoG Subject 2 did not make manual responses because these responses were found to interfere with visual fixation. **B)** ECoG responses were measured in an event related design, where stimuli where presented every 1000 ms. Stimuli were presented for 500 ms followed by a blank screen. **C)** Stimuli were presented in blocks of 12 seconds during fMRI, followed by 12 seconds of blank. **D)** Example ECoG power spectrum for one electrode. ECoG data showed broadband increases (>100Hz) compared to baseline, narrowband gamma increases around 40 HZ, and a decrease in alpha power around 10 Hz. **E)** The BOLD response increased in different levels for the different stimuli averaged across subjects. When averaging the BOLD signal across subjects, the percent signal change per subjects was vector length normalized, 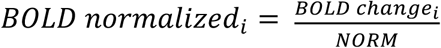, for condition *i*, in 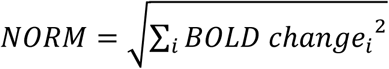. To then re-estimate the percent signal change across subjects, the averaged vector length normalized values were multiplied by the average norm.

**Figure S3.**
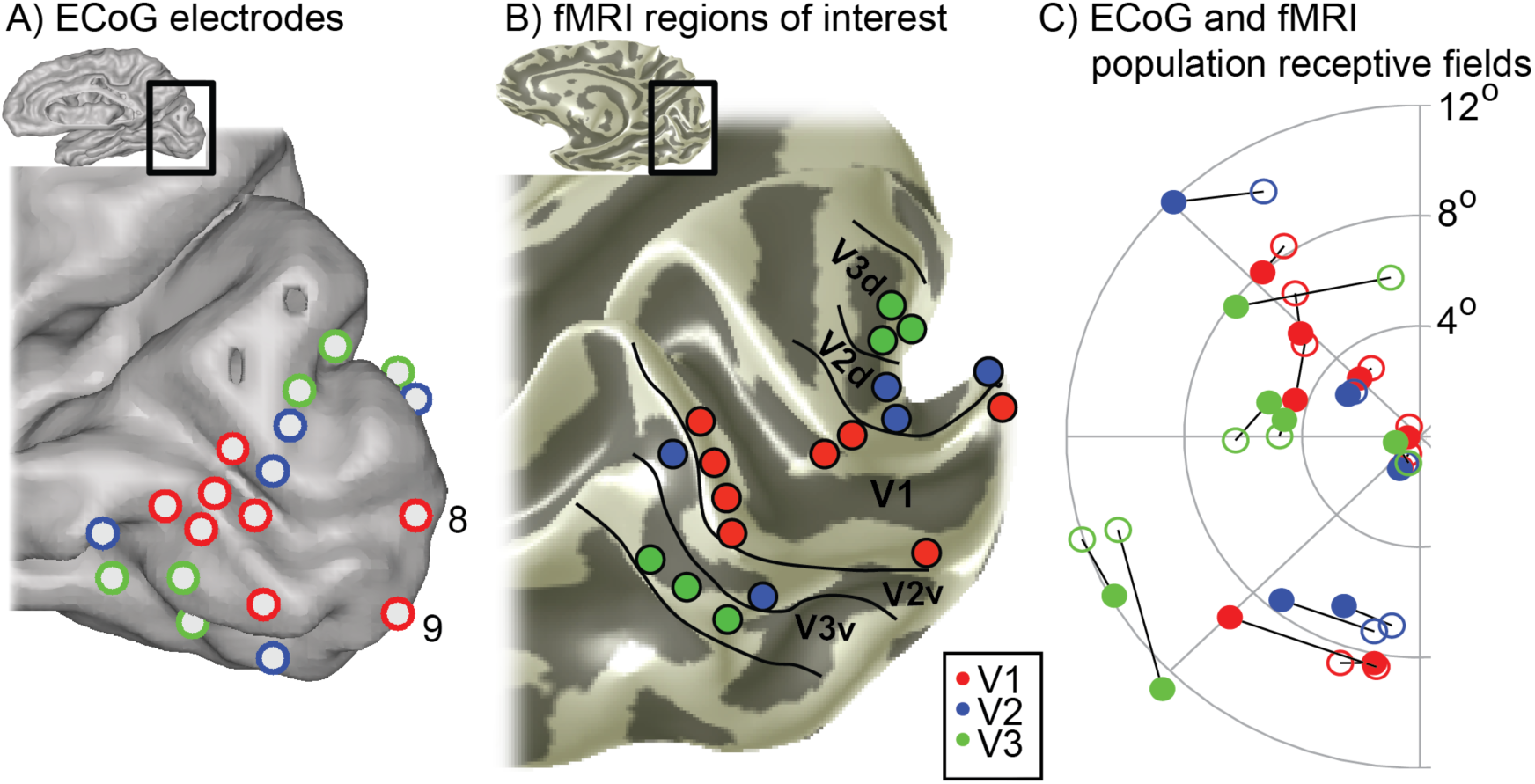
ROI selection. **A)** Channels in ECoG S1 selected for further analysis. These channels were located within V1 (red), V2 (blue), or V3 (green), had significant broadband or gamma response to any stimuli, and had pRF variance explained >0.15. V1 sites 8 and 9 are indicated, since these had the largest gamma responses. **B)** Electrode ROIs in fMRI S1. Disc ROIs (radius = 2 mm) were defined to have similar anatomical and retinotopic position as the ECoG Channels. **C)** The pRF centers for fMRI ROIs (filled circles) were chosen to be close to those for ECoG electrodes (open circles). Because the pRF centers measured with fMRI do not completely cover the visual field map, the locations can differ slightly.

**Figure S4.**
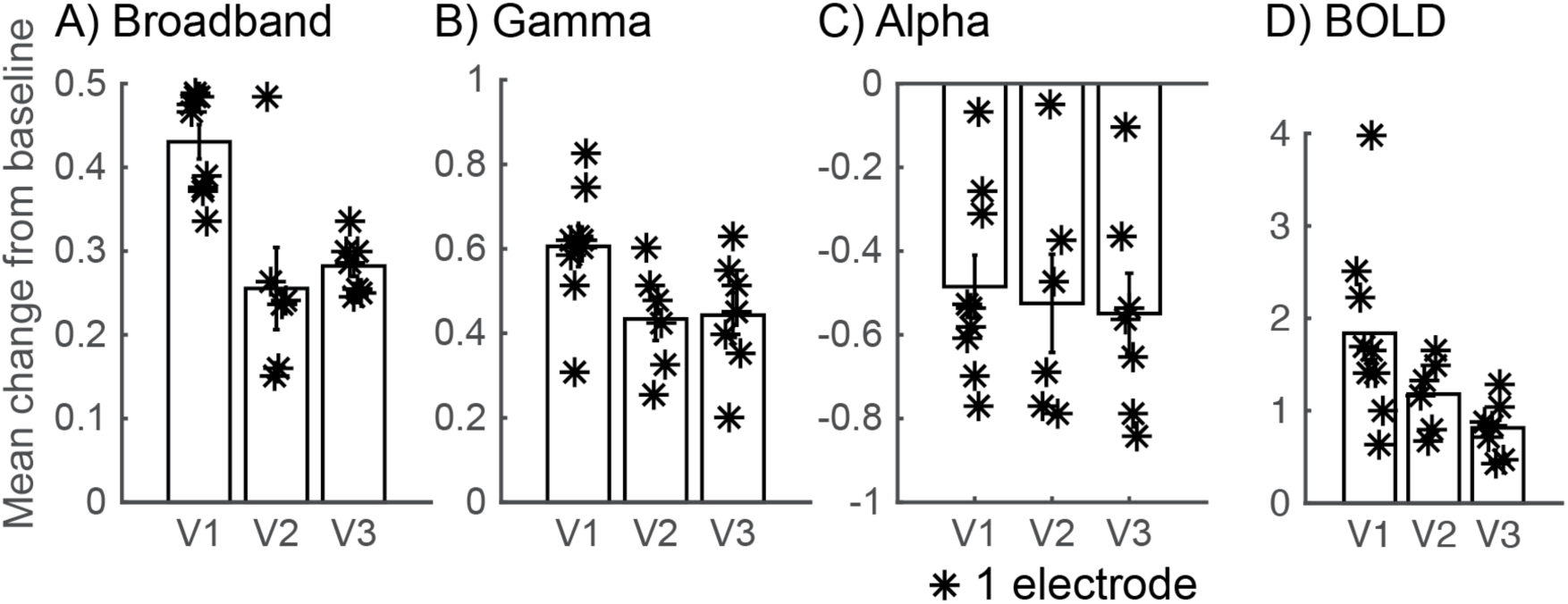
Broadband, gamma and alpha changes in V1, V2 and V3. **(A)** For each ECoG electrode, for each stimulus condition, the broadband change was calculated. The average log10 power from the inter stimulus baseline period was subtracted. The mean change from the baseline was then averaged across the 8-10 stimuli. **(B and C)** The same as A) shown for gamma and alpha. **(D)** For each ECoG electrode, for each stimulus condition, the BOLD percent signal change was calculated. The mean change from the baseline was then averaged across the stimuli.

**Figure S5.**
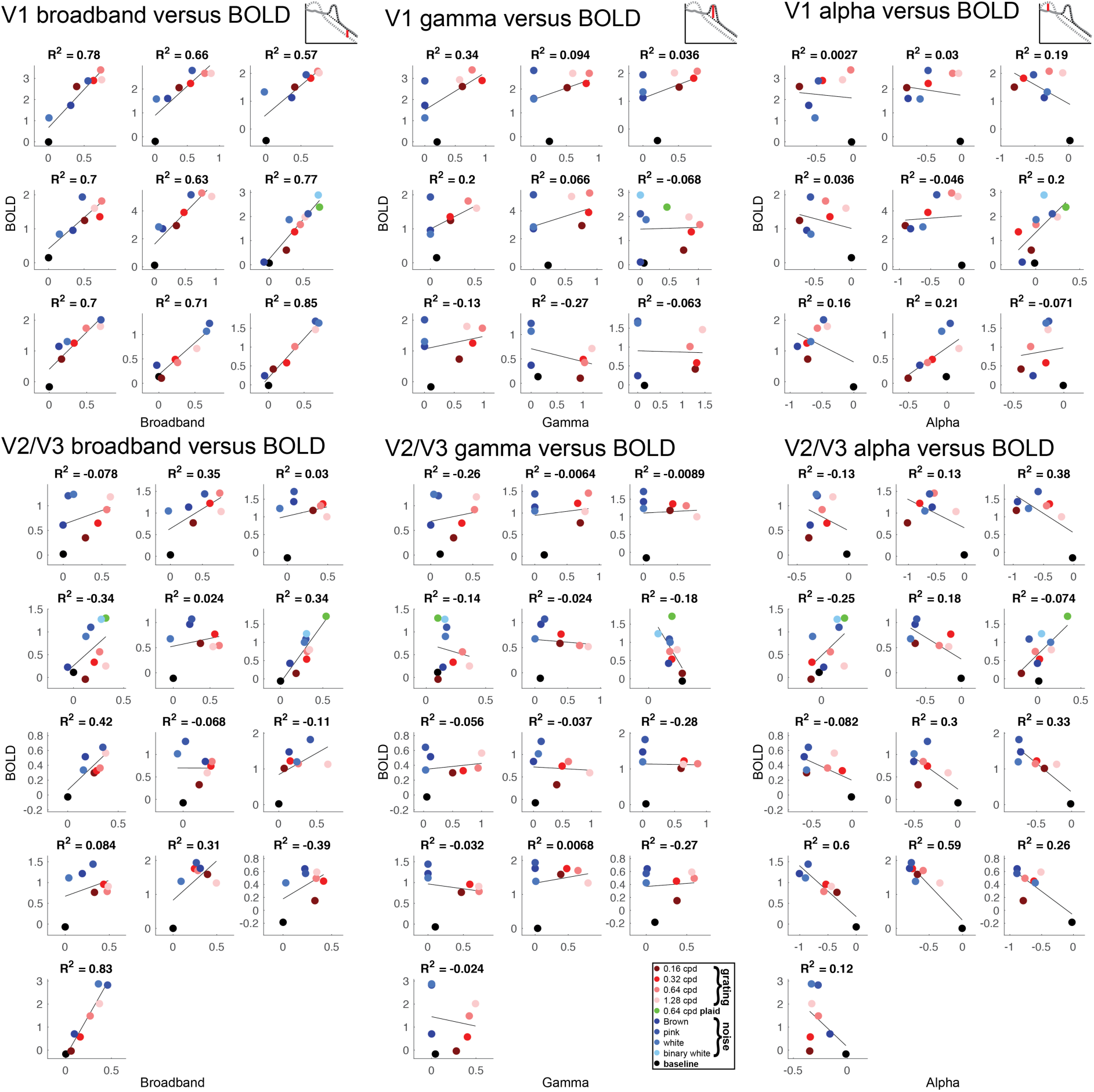
Correlation between BOLD and ECoG broadband, gamma and alpha for all electrodes. Correlation between BOLD and ECoG in V1 and V2/V3. The R^2^ is cross-validated: beta values are calculated from half the ECoG trials and half the fMRI subjects, and the regression model is tested on the other half of the trials and subjects.

**Figure S6.**
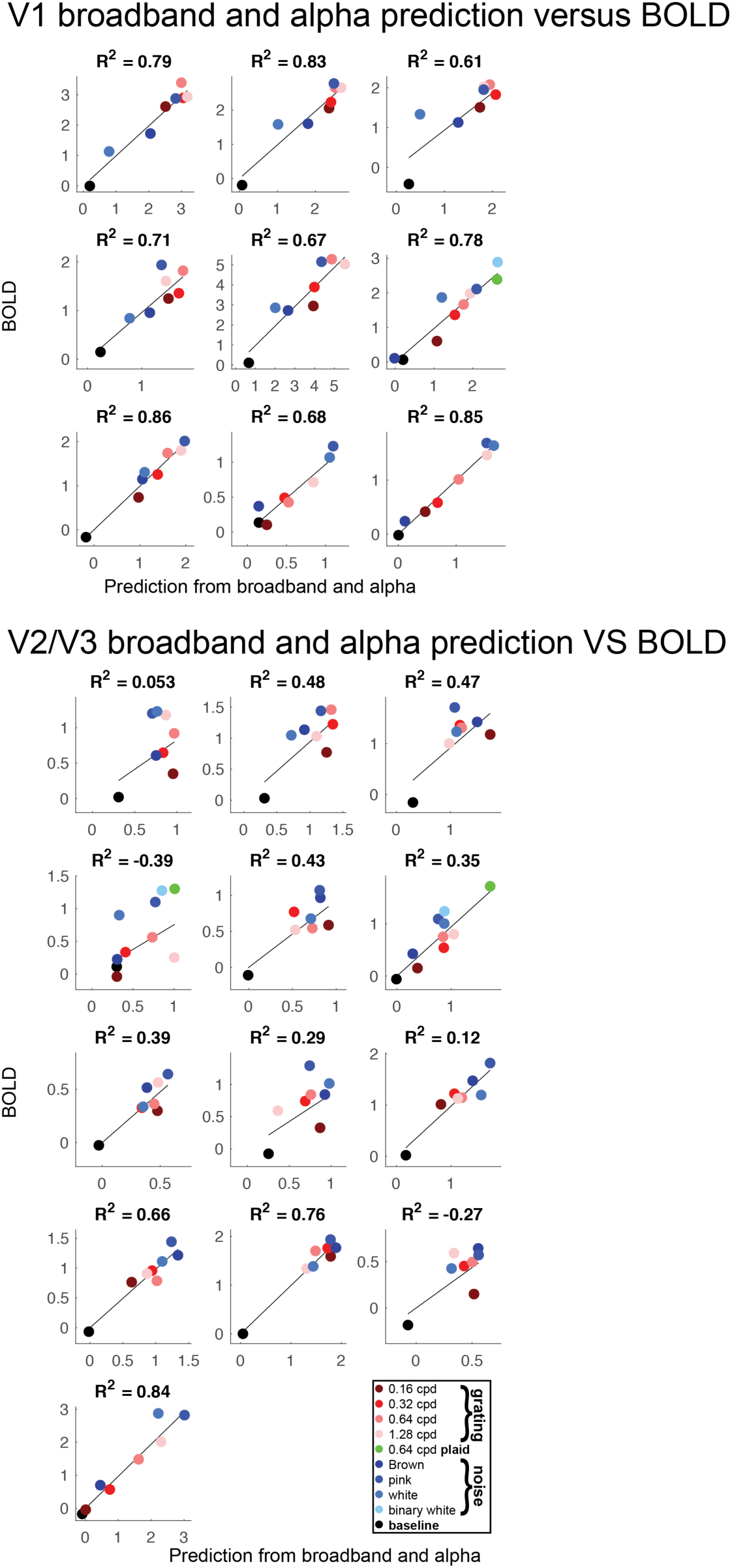
BOLD predicted by ECoG broadband and alpha for all electrodes. This plots shows the predicted BOLD (x-axis) versus measured BOLD (y-axis) for the 9 V1 sites (top) and 13 V2/V3 sites (bottom), based on a linear regression of the broadband and alpha components of the ECoG signals. The coefficient of determination, R^2^, was cross-validated.

**Figure S7:**
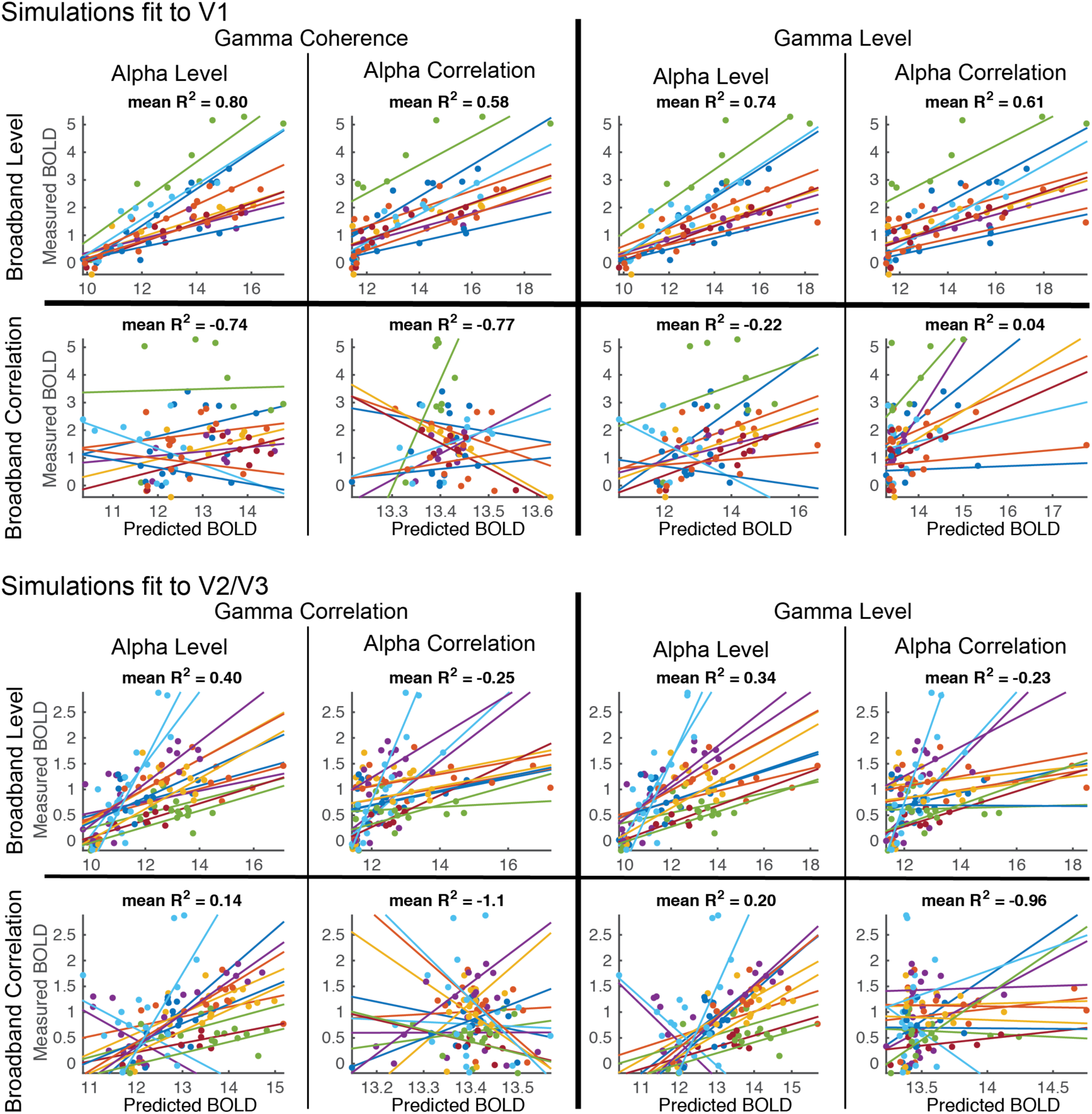
Accuracy of BOLD predictions from simulated neuronal activity. This plots shows the predicted BOLD (x-axis) versus measured BOLD (y-axis) for the 9 V1 sites (top) and 13 V2/V3 sites (bottom). Each color corresponds to one site. The cross-validated coefficient of determination (R^2^) was computed separately for each of the 9 sites, and then averaged. The different subplots are models solved with different constraints. In the main text of the paper, model parameters were fit with three constraints: (1) the C^1^ (broadband) time series had a fixed, non-zero level (but could vary in correlation between neurons), (2) the C^2^ (gamma) time series had a fixed, non-zero level (but could vary in correlation), and (3) the C^3^ time series had a fixed, non-zero correlation (but could vary in level). The model predictions based on these constrains are plotted in the upper left of both the upper panel (V1) and the lower panel (V2/V3). Seven alternative models were run, and their predictions are shown in the remaining panels. For these models, the three input types, C^1^, C^2^, and C^3^, were constrained to have time series varying in either the level or correlations across neurons, but not both.

**Figure S8.**
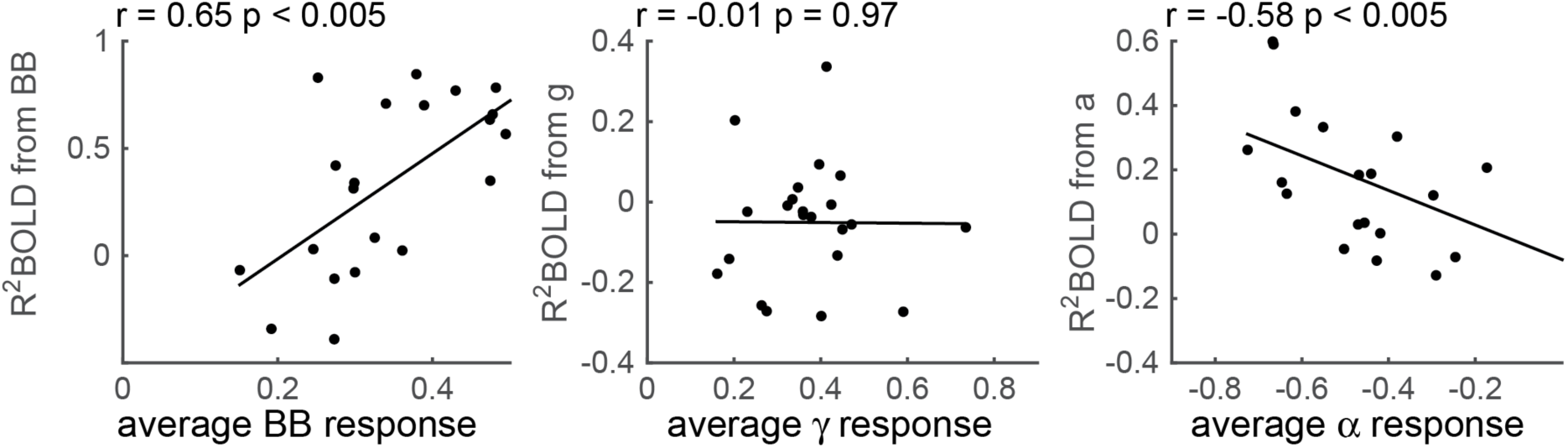
Variance in the BOLD response explained by ECoG (R^2^, the coefficient of determination) as a function of the size of the ECoG response. Each dot represents one electrode. X-axis: for each electrode, ECoG broadband, gamma and alpha responses were averaged across (non-baseline) stimuli. Y-axis: the cross-validated R^2^ when BOLD is explained by broadband (left), gamma (middle) and alpha (right).

**Figure S9.**
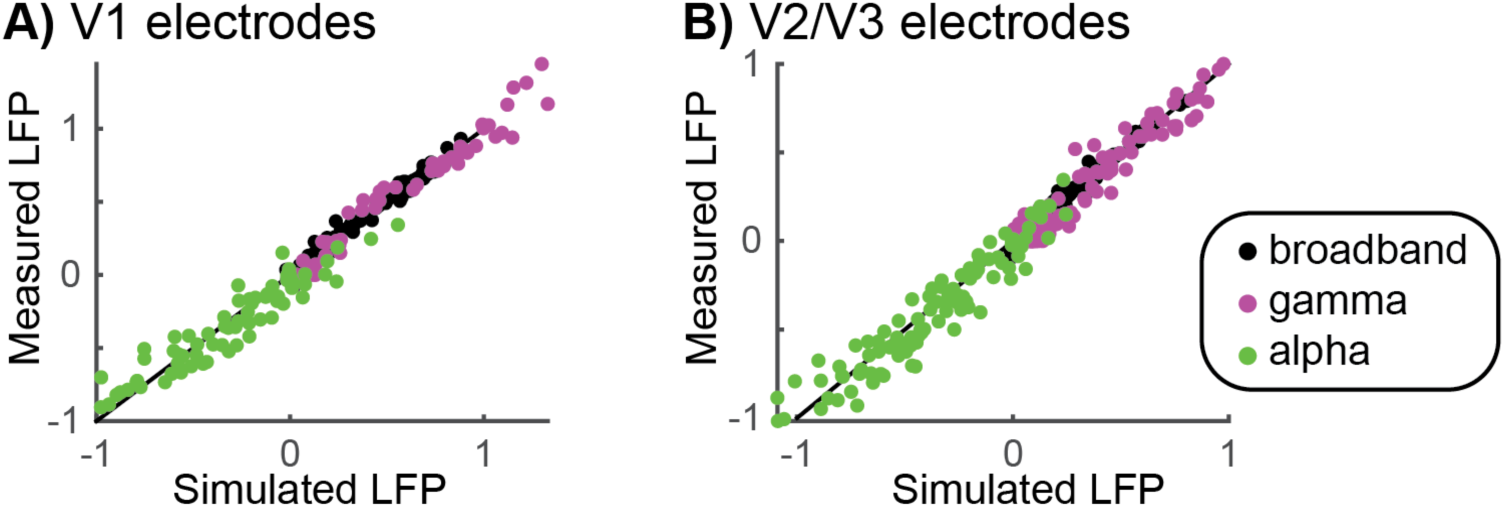
Relation between the simulated and measured LFP values. Every dot represents the broadband (black), gamma (magenta) or alpha (green) power change for one electrode, one stimulus condition. The power changes in the LFP are driven by changes in parameters *C^1^*, *C^2^* and *C^3^*. We fitted these parameters such that the simulated LFP values for broadband, gamma and alpha nicely match the measured values.

## References

1. Logothetis NK, Wandell BA (2004) Interpreting the BOLD signal. Annual review of physiology 66: 735–769.

2. Buzsaki G, Anastassiou CA, Koch C (2012) The origin of extracellular fields and currents--EEG, ECoG, LFP and spikes. Nat Rev Neurosci 13: 407–420.

3. Hämäläinen M, Hari R, Ilmoniemi RJ, Knuutila J, Lounasmaa OV (1993) Magnetoencephalography—theory, instrumentation, and applications to noninvasive studies of the working human brain. Reviews of modern Physics 65: 413.

4. George JS, Aine CJ, Mosher JC, Schmidt DM, Ranken DM, et al. (1995) Mapping function in the human brain with magnetoencephalography, anatomical magnetic resonance imaging, and functional magnetic resonance imaging. J Clin Neurophysiol 12: 406–431.

5. Cottereau BR, Ales JM, Norcia AM (2015) How to use fMRI functional localizers to improve EEG/MEG source estimation. J Neurosci Methods 250: 64–73.

6. Hermes D, Miller KJ, Noordmans HJ, Vansteensel MJ, Ramsey NF (2010) Automated electrocorticographic electrode localization on individually rendered brain surfaces. Journal of neuroscience methods 185: 293–298.

7. Huk AC, Ress D, Heeger DJ (2001) Neuronal basis of the motion aftereffect reconsidered. Neuron 32: 161–172.

8. Ales JM, Norcia AM (2009) Assessing direction-specific adaptation using the steady-state visual evoked potential: results from EEG source imaging. J Vis 9: 8.

9. Pestilli F, Carrasco M, Heeger DJ, Gardner JL (2011) Attentional enhancement via selection and pooling of early sensory responses in human visual cortex. Neuron 72: 832–846.

10. Itthipuripat S, Ester EF, Deering S, Serences JT (2014) Sensory gain outperforms efficient readout mechanisms in predicting attention-related improvements in behavior. The Journal of neuroscience : the official journal of the Society for Neuroscience 34: 13384–13398.

11. Winawer J, Kay KN, Foster BL, Rauschecker AM, Parvizi J, et al. (2013) Asynchronous broadband signals are the principal source of the BOLD response in human visual cortex. Curr Biol 23: 1145–1153.

12. Goense JB, Logothetis NK (2008) Neurophysiology of the BOLD fMRI signal in awake monkeys. Current biology : CB 18: 631–640.

13. Niessing J, Ebisch B, Schmidt KE, Niessing M, Singer W, et al. (2005) Hemodynamic signals correlate tightly with synchronized gamma oscillations. Science 309: 948–951.

14. Mukamel R, Gelbard H, Arieli A, Hasson U, Fried I, et al. (2005) Coupling between neuronal firing, field potentials, and FMRI in human auditory cortex. Science 309: 951–954.

15. Shmuel A, Augath M, Oeltermann A, Logothetis NK (2006) Negative functional MRI response correlates with decreases in neuronal activity in monkey visual area V1. Nature neuroscience 9: 569–577.

16. Logothetis NK, Pauls J, Augath M, Trinath T, Oeltermann A (2001) Neurophysiological investigation of the basis of the fMRI signal. Nature 412: 150–157.

17. Lachaux JP, Fonlupt P, Kahane P, Minotti L, Hoffmann D, et al. (2007) Relationship between task-related gamma oscillations and BOLD signal: new insights from combined fMRI and intracranial EEG. Hum Brain Mapp 28: 1368–1375.

18. Muthukumaraswamy SD, Singh KD (2009) Functional decoupling of BOLD and gamma-band amplitudes in human primary visual cortex. Hum Brain Mapp 30: 2000–2007.

19. Swettenham JB, Muthukumaraswamy SD, Singh KD (2013) BOLD Responses in Human Primary Visual Cortex are Insensitive to Substantial Changes in Neural Activity. Frontiers in human neuroscience 7: 76.

20. Maier A, Wilke M, Aura C, Zhu C, Ye FQ, et al. (2008) Divergence of fMRI and neural signals in V1 during perceptual suppression in the awake monkey. Nature neuroscience 11: 1193–1200.

21. Scheeringa R, Fries P, Petersson KM, Oostenveld R, Grothe I, et al. (2011) Neuronal dynamics underlying high-and low-frequency EEG oscillations contribute independently to the human BOLD signal. Neuron 69: 572–583.

22. Harvey BM, Vansteensel MJ, Ferrier CH, Petridou N, Zuiderbaan W, et al. (2013) Frequency specific spatial interactions in human electrocorticography: V1 alpha oscillations reflect surround suppression. NeuroImage 65: 424–432.

23. Scheeringa R, Koopmans PJ, van Mourik T, Jensen O, Norris DG (2016) The relationship between oscillatory EEG activity and the laminar-specific BOLD signal. Proceedings of the National Academy of Sciences of the United States of America.

24. Lima B, Cardoso MM, Sirotin YB, Das A (2014) Stimulus-related neuroimaging in task-engaged subjects is best predicted by concurrent spiking. The Journal of neuroscience : the official journal of the Society for Neuroscience 34: 13878–13891.

25. Rees G, Friston K, Koch C (2000) A direct quantitative relationship between the functional properties of human and macaque V5. Nat Neurosci 3: 716–723.

26. Heeger DJ, Huk AC, Geisler WS, Albrecht DG (2000) Spikes versus BOLD: what does neuroimaging tell us about neuronal activity? Nat Neurosci 3: 631–633.

27. Heeger DJ, Ress D (2002) What does fMRI tell us about neuronal activity? Nat Rev Neurosci 3: 142–151.

28. Mathiesen C, Caesar K, Akgoren N, Lauritzen M (1998) Modification of activity-dependent increases of cerebral blood flow by excitatory synaptic activity and spikes in rat cerebellar cortex. J Physiol 512 (Pt 2): 555–566.

29. Lee JH, Durand R, Gradinaru V, Zhang F, Goshen I, et al. (2010) Global and local fMRI signals driven by neurons defined optogenetically by type and wiring. Nature 465: 788–792.

30. Zanos TP, Mineault PJ, Pack CC (2011) Removal of spurious correlations between spikes and local field potentials. J Neurophysiol 105: 474–486.

31. Pesaran B (2008) Spectral analysis for neural signals. In: Mitra P, editor. Neural Signal Processing: Quantitative Analysis of Neural Activity: Society for Neuroscience. pp. 1–12.

32. Raichle ME, Mintun MA (2006) Brain work and brain imaging. Annu Rev Neurosci 29: 449–476.

33. Yu BM, Cunningham JP, Santhanam G, Ryu SI, Shenoy KV, et al. (2009) Gaussian-process factor analysis for low-dimensional single-trial analysis of neural population activity. J Neurophysiol 102: 614–635.

34. Stevenson IH, Kording KP (2011) How advances in neural recording affect data analysis. Nat Neurosci 14: 139–142.

35. Hermes D, Miller KJ, Wandell BA, Winawer J (2015) Stimulus dependence of gamma oscillations in human visual cortex. Cerebral cortex 25: 2951–2959.

36. Koch C, Rapp M, Segev I (1996) A brief history of time (constants). Cereb Cortex 6: 93–101.

37. Miller KJ, Sorensen LB, Ojemann JG, den Nijs M (2009) Power-law scaling in the brain surface electric potential. PLoS computational biology 5: e1000609.

38. Henrie JA, Shapley R (2005) LFP power spectra in V1 cortex: the graded effect of stimulus contrast. J Neurophysiol 94: 479–490.

39. Tan AY, Chen Y, Scholl B, Seidemann E, Priebe NJ (2014) Sensory stimulation shifts visual cortex from synchronous to asynchronous states. Nature 509: 226–229.

40. Jia X, Tanabe S, Kohn A (2013) gamma and the coordination of spiking activity in early visual cortex. Neuron 77: 762–774.

41. Hasenstaub A, Shu Y, Haider B, Kraushaar U, Duque A, et al. (2005) Inhibitory postsynaptic potentials carry synchronized frequency information in active cortical networks. Neuron 47: 423–435.

42. Gray CM, Konig P, Engel AK, Singer W (1989) Oscillatory responses in cat visual cortex exhibit inter-columnar synchronization which reflects global stimulus properties. Nature 338: 334–337.

43. Gray CM, Singer W (1989) Stimulus-specific neuronal oscillations in orientation columns of cat visual cortex. Proceedings of the National Academy of Sciences of the United States of America 86: 1698–1702.

44. Perrenoud Q, Pennartz CM, Gentet LJ (2016) Membrane Potential Dynamics of Spontaneous and Visually Evoked Gamma Activity in V1 of Awake Mice. PLoS biology 14: e1002383.

45. Burns SP, Xing D, Shapley RM (2011) Is gamma-band activity in the local field potential of V1 cortex a “clock” or filtered noise? The Journal of neuroscience : the official journal of the Society for Neuroscience 31: 9658–9664.

46. Jensen O, Mazaheri A (2010) Shaping functional architecture by oscillatory alpha activity: gating by inhibition. Front Hum Neurosci 4: 186.

47. Schalk G (2015) A general framework for dynamic cortical function: the function-throughbiased-oscillations (FBO) hypothesis. Frontiers in human neuroscience 9: 352.

48. Llinas RR, Grace AA, Yarom Y (1991) In vitro neurons in mammalian cortical layer 4 exhibit intrinsic oscillatory activity in the 10-to 50-Hz frequency range. Proc Natl Acad Sci U S A 88: 897–901.

49. Ojemann GA, Ojemann J, Ramsey NF (2013) Relation between functional magnetic resonance imaging (fMRI) and single neuron, local field potential (LFP) and electrocorticography (ECoG) activity in human cortex. Frontiers in human neuroscience 7: 34.

50. Hermes D, Miller KJ, Wandell BA, Winawer J (2015) Gamma oscillations in visual cortex: the stimulus matters. Trends Cogn Sci 19: 57–58.

51. Ray S, Maunsell JH (2011) Different origins of gamma rhythm and high-gamma activity in macaque visual cortex. PLoS biology 9: e1000610.

52. Singer W (1999) Neuronal synchrony: a versatile code for the definition of relations? Neuron 24: 49-65, 111-125.

53. Buzsaki G, Wang XJ (2012) Mechanisms of gamma oscillations. Annu Rev Neurosci 35: 203–225.

54. Miller KJ, Zanos S, Fetz EE, den Nijs M, Ojemann JG (2009) Decoupling the cortical power spectrum reveals real-time representation of individual finger movements in humans. The Journal of neuroscience : the official journal of the Society for Neuroscience 29: 3132–3137.

55. Manning JR, Jacobs J, Fried I, Kahana MJ (2009) Broadband shifts in local field potential power spectra are correlated with single-neuron spiking in humans. The Journal of neuroscience : the official journal of the Society for Neuroscience 29: 13613–13620.

56. Musall S, von Pfostl V, Rauch A, Logothetis NK, Whittingstall K (2014) Effects of neural synchrony on surface EEG. Cereb Cortex 24: 1045–1053.

57. Engell AD, Huettel S, McCarthy G (2012) The fMRI BOLD signal tracks electrophysiological spectral perturbations, not event-related potentials. NeuroImage 59: 2600–2606.

58. Sirotin YB, Das A (2009) Anticipatory haemodynamic signals in sensory cortex not predicted by local neuronal activity. Nature 457: 475–479.

59. Burns SP, Xing D, Shapley RM (2010) Comparisons of the dynamics of local field potential and multiunit activity signals in macaque visual cortex. The Journal of neuroscience : the official journal of the Society for Neuroscience 30: 13739–13749.

60. Jia X, Xing D, Kohn A (2013) No consistent relationship between gamma power and peak frequency in macaque primary visual cortex. The Journal of neuroscience : the official journal of the Society for Neuroscience 33: 17–25.

61. Dalal SS, Baillet S, Adam C, Ducorps A, Schwartz D, et al. (2009) Simultaneous MEG and intracranial EEG recordings during attentive reading. NeuroImage 45: 1289–1304.

62. Adrian ED, Matthews BHC (1934) The Berger rhythm: Potential changes from the occipital lobes in man. Brain 57: 355–385.

63. Norcia AM, Appelbaum LG, Ales JM, Cottereau BR, Rossion B (2015) The steady-state visual evoked potential in vision research: A review. J Vis 15: 4.

64. Tallon-Baudry C, Bertrand O (1999) Oscillatory gamma activity in humans and its role in object representation. Trends Cogn Sci 3: 151–162.

65. Hermes D, Miller KJ, Vansteensel MJ, Aarnoutse EJ, Leijten FS, et al. (2012) Neurophysiologic correlates of fMRI in human motor cortex. Human brain mapping 33: 1689–1699.

66. Hermes D, Miller KJ, Vansteensel MJ, Edwards E, Ferrier CH, et al. (2014) Cortical theta wanes for language. NeuroImage 85 Pt 2: 738–748.

67. Magri C, Schridde U, Murayama Y, Panzeri S, Logothetis NK (2012) The amplitude and timing of the BOLD signal reflects the relationship between local field potential power at different frequencies. The Journal of neuroscience : the official journal of the Society for Neuroscience 32: 1395–1407.

68. Harris KD, Thiele A (2011) Cortical state and attention. Nat Rev Neurosci 12: 509–523.

69. Mazaheri A, Jensen O (2010) Rhythmic pulsing: linking ongoing brain activity with evoked responses. Front Hum Neurosci 4: 177.

70. Mazaheri A, Jensen O (2008) Asymmetric amplitude modulations of brain oscillations generate slow evoked responses. J Neurosci 28: 7781–7787.

71. Raichle ME (2010) Two views of brain function. Trends in cognitive sciences 14: 180–190.

72. Samaha J, Gosseries O, Postle BR (2017) Distinct Oscillatory Frequencies Underlie Excitability of Human Occipital and Parietal Cortex. J Neurosci 37: 2824–2833.

73. Dugue L, Marque P, VanRullen R (2011) The phase of ongoing oscillations mediates the causal relation between brain excitation and visual perception. J Neurosci 31: 11889–11893.

74. Scheeringa R, Mazaheri A, Bojak I, Norris DG, Kleinschmidt A (2011) Modulation of visually evoked cortical FMRI responses by phase of ongoing occipital alpha oscillations. J Neurosci 31: 3813–3820.

75. Kang K, Shelley M, Henrie JA, Shapley R (2010) LFP spectral peaks in V1 cortex: network resonance and cortico-cortical feedback. Journal of computational neuroscience 29: 495–507.

76. Xing D, Shen Y, Burns S, Yeh CI, Shapley R, et al. (2012) Stochastic generation of gamma-band activity in primary visual cortex of awake and anesthetized monkeys. J Neurosci 32: 13873–13880a.

77. Sloan HL, Austin VC, Blamire AM, Schnupp JW, Lowe AS, et al. (2010) Regional differences in neurovascular coupling in rat brain as determined by fMRI and electrophysiology. NeuroImage 53: 399–411.

78. Conner CR, Ellmore TM, Pieters TA, DiSano MA, Tandon N (2011) Variability of the relationship between electrophysiology and BOLD-fMRI across cortical regions in humans. The Journal of neuroscience : the official journal of the Society for Neuroscience 31: 12855–12865.

79. Huo BX, Smith JB, Drew PJ (2014) Neurovascular coupling and decoupling in the cortex during voluntary locomotion. The Journal of neuroscience : the official journal of the Society for Neuroscience 34: 10975–10981.

80. Goense J, Merkle H, Logothetis NK (2012) High-resolution fMRI reveals laminar differences in neurovascular coupling between positive and negative BOLD responses. Neuron 76: 629–639.

81. Zaldivar D, Rauch A, Whittingstall K, Logothetis NK, Goense J (2014) Dopamine-induced dissociation of BOLD and neural activity in macaque visual cortex. Curr Biol 24: 2805–2811.

82. Heeger DJ, Simoncelli EP, Movshon JA (1996) Computational models of cortical visual processing. Proc Natl Acad Sci U S A 93: 623–627.

83. Carandini M, Heeger DJ (2012) Normalization as a canonical neural computation. Nat Rev Neurosci 13: 51–62.

84. Kay KN, Winawer J, Rokem A, Mezer A, Wandell BA (2013) A two-stage cascade model of BOLD responses in human visual cortex. PLoS computational biology 9: e1003079.

85. Wandell BA (1995) Foundations of vision. Sunderland, Mass.: Sinauer Associates. xvi, 476 p., [474] p. of plates p.

86. Carandini M, Demb JB, Mante V, Tolhurst DJ, Dan Y, et al. (2005) Do we know what the early visual system does? J Neurosci 25: 10577–10597.

87. Olshausen BA, Field DJ (2005) How close are we to understanding v1? Neural Comput 17: 1665–1699.

88. Gavish M, Donoho D (2012) Three Dream Applications of Verifiable Computational Results. Computing in Science & Engineering 14: 26–31.

89. LeVeque RJ, Mitchell IM, Stodden V (2012) Reproducible Research for Scientific Computing: Tools and Strategies for Changing the Culture. Computing in Science & Engineering 14: 13–17.

90. Axelrod V (2014) Minimizing bugs in cognitive neuroscience programming. Frontiers in Psychology 5.

91. Bedard C, Destexhe A (2009) Macroscopic models of local field potentials and the apparent 1/f noise in brain activity. Biophys J 96: 2589–2603.

92. Milstein J, Mormann F, Fried I, Koch C (2009) Neuronal shot noise and Brownian 1/f2 behavior in the local field potential. PLoS one 4: e4338.

93. Bedard C, Kroger H, Destexhe A (2006) Does the 1/f frequency scaling of brain signals reflect self-organized critical states? Phys Rev Lett 97: 118102.

94. Logothetis NK, Kayser C, Oeltermann A (2007) In vivo measurement of cortical impedance spectrum in monkeys: implications for signal propagation. Neuron 55: 809–823.

95. Jia X, Smith MA, Kohn A (2011) Stimulus selectivity and spatial coherence of gamma components of the local field potential. The Journal of neuroscience : the official journal of the Society for Neuroscience 31: 9390–9403.

96. Welch PD (1967) Use of Fast Fourier Transform for Estimation of Power Spectra - a Method Based on Time Averaging over Short Modified Periodograms. Ieee Transactions on Audio and Electroacoustics Au15: 70-&.

97. Benson NC, Butt OH, Datta R, Radoeva PD, Brainard DH, et al. (2012) The retinotopic organization of striate cortex is well predicted by surface topology. Curr Biol 22: 2081–2085.

98. Smith AM, Lewis BK, Ruttimann UE, Ye FQ, Sinnwell TM, et al. (1999) Investigation of low frequency drift in fMRI signal. NeuroImage 9: 526–533.

99. Kay KN, Rokem A, Winawer J, Dougherty RF, Wandell BA (2013) GLMdenoise: a fast, automated technique for denoising task-based fMRI data. Front Neurosci 7: 247.

100. Dumoulin SO, Wandell BA (2008) Population receptive field estimates in human visual cortex. NeuroImage 39: 647–660.

101. Winawer J, Horiguchi H, Sayres RA, Amano K, Wandell BA (2010) Mapping hV4 and ventral occipital cortex: the venous eclipse. Journal of vision 10: 1.

